# Social isolation uncovers a brain-wide circuit underlying context-dependent territory-covering micturition behavior

**DOI:** 10.1101/798132

**Authors:** Minsuk Hyun, Julian Taranda, Gianna Radeljic, Lauren Miner, Wengang Wang, Nicole Ochandarena, Kee Wui Huang, Pavel Osten, Bernardo Sabatini

**Affiliations:** Howard Hughes Medical Institute, Department of Neurobiology, Harvard Medical School, Boston, Massachusetts, 02115, USA; Cold Spring Harbor Laboratory, Cold Spring Harbor, New York, 11724, USA

## Abstract

The controlled and volitional release of urine, or micturition, serves a fundamental physiological function and, in many species, is critical for social communication. In mice, the decision to release urine is modulated by external and internal factors such as environmental stimuli and social history and is transmitted to the spinal cord via the pontine micturition center (PMC). The neural pathways by which social experience and sensory stimuli interact to control PMC activity and regulate micturition are unclear. Here we establish a behavioral paradigm in which mice, depending on their strain, social experience, and immediate sensory context, display either high or low territory-covering micturition (TCM). We demonstrate that social context is represented by coordinated global activity changes in the urination network upstream of the PMC, whereas sensory context is represented by the activation of discrete nodes in the network. Furthermore, we show that the lateral hypothalamic area (LHA), which is directly upstream of PMC, is a key node that can switch micturition behavior between high and low TCM modes.

## Introduction

The control of urine output, or micturition, is necessary for the survival of animals in the wild. Mice, like many mammals, innately modulate micturition in response to internal and environmental cues and alter the spatiotemporal patterns of urine output depending on their strain, sex, and position in the social hierarchy. For example, when male mice detect competitors or potential mates (Hurst and Beynon, 2004), they assiduously deposit urine throughout the environment, a process often referred to ‘territorial marking’ or ‘scent marking’. Social experience further modulates the extent and pattern of such urine marking by adult males: “dominant” mice will vigorously urinate with small and dispersed urine spots whereas “subordinate” males limit urination to large, individual spots, often located in the corners of the territory (Desjardins et al., 1973; Hou et al., 2016). Therefore, the decision to release urine results from the complex integration and processing of many signals, including sensory inputs, past social experience, and the state of the bladder. However, unlike many other complex decision processes, micturition is carried out by the coordinated activity of only three muscles – contraction of the bladder wall detrusor muscle and relaxation of the internal and external urethral sphincters (Groat and Wickens, 2013).

The brainstem pontine micturition center (PMC), also eponymously referred to as Barrington’s nucleus, is the output nucleus of the brain that controls the bladder and, therefore, serves as the command center of the brain to regulate micturition (Drake et al., 2010; Groat and Wickens, 2013; Hou et al., 2016; Valentino et al., 1994; Yao et al., 2018). Recent studies have begun to shed light on how different cell-types in the PMC work together to coordinate bladder output. Glutamatergic PMC neurons marked by expression of Corticotropin-releasing hormone (*Crh*, also known as Corticotropin-releasing factor) bilaterally innervate the sacral spinal cord where motor neurons innervating the bladder wall reside (Hou et al., 2016). In parallel, a recent study utilizing *Esr1*^ires-Cre^ mouse line demonstrated a potentially distinct glutamatergic population that controls the urethral sphincters to permit urine release (Keller et al., 2018).

The PMC, including both the *Crh*-expressing and sphincter-controlling neurons, receives inputs from distributed circuits across the brain which presumably carry pro- and anti-micturition information that is processed in the PMC (Hou et al., 2016; Verstegen et al., 2019; Yao et al., 2018). Thus, PMC serves as a suitable starting point to retrogradely map the neural circuitry by which information is integrated to trigger a specific motor action. Nevertheless, we do not know when different inputs to PMC are active during urinary marking behavior, how these are altered by olfactory cues, and how they modulate the activity of PMC.

In order to address these unknowns, we designed a behavioral paradigm to specifically study “territory-covering micturition” (TCM) in which a male mouse exposed to female urine robustly covers the arena with urine. Here, we refer to this behavior as TCM since it is descriptive of the action and does not imply a specific intent or goal of the mouse. We find that the propensity of each mouse to display TCM depends on its strain, social dominance, olfactory context, and previous experience in the behavioral arena. Socially-housed male C57BL6/J mice do not display robust TCM, but this behavior can be induced by social isolation. Mapping of brain-wide inputs to *Crh+* PMC neurons suggest that the connectivity upstream of these neurons is not affected by social isolation. In contrast, brain-wide mapping of the immediate early gene c-fos mapping revealed that the activities of upstream areas differ in socially-isolated and group housed mice exposed to urine. Comparing the whole-brain activity of isolated males with high TCM and low TCM we identified putative key nodes of the upstream PMC micturition network that are capable of carrying diverse TCM-regulating signals to the bladder. Furthermore, chemogenetic modulation of one of the identified nodes of the putative PMC micturition network – the lateral hypothalamus – bidirectionally modulates TCM. Thus, social-environmental context is represented by coordinated activity changes in the upstream PMC micturition network, whereas discrete nodes in the upstream PMC micturition network, including LHA, represent TCM-relevant sensory context. These findings provide new insights into brain-wide mechanisms governing TCM and motivate next steps to fully trace the sensorimotor transformation underlying innate TCM.

## Results

### Social isolation reveals extensive territory-covering micturition

Inbred laboratory mice of different strains are genetically and behaviorally diverse (Crawley et al., 1997) and therefore may display differences in micturition behaviors. To systematically characterize such potential differences, we examined the TCM patterns of 4 commonly used inbred strains: C57BL6/J, BALBc, 129s, CD1. Pair-housed adult males of different strains were placed into cages individually, with 50 µl of male urine placed in the center and filter paper lining the floor. After 1 hour in the cage, the filter paper and mouse were removed and the pattern of TCM was analyzed post-hoc under blue LED illumination (Fig. 1a). The most commonly used strain, C57BL6/J male mice, displayed the lowest TCM whereas BALBc male mice displayed highest TCM (Fig. 1b, C57BL6/J: 18 ± 7 marks; BALBC: marks 132 ± 39 marks; n=8 mice per strain). Inspection of urine marks revealed a clear separation in TCM in each partner, such that one mouse displayed “high TCM” and the other displayed “low TCM” (Supplementary Fig. 1a). This pattern of differential micturition within a pair of male mice was previously associated with known factors such as social rank (Desjardins et al., 1973; Hou et al., 2016). Thus, the extent of TCM and its regulation through social experience are dependent on the strain, and therefore genetics, of individual male mice.

**Figure 1.**
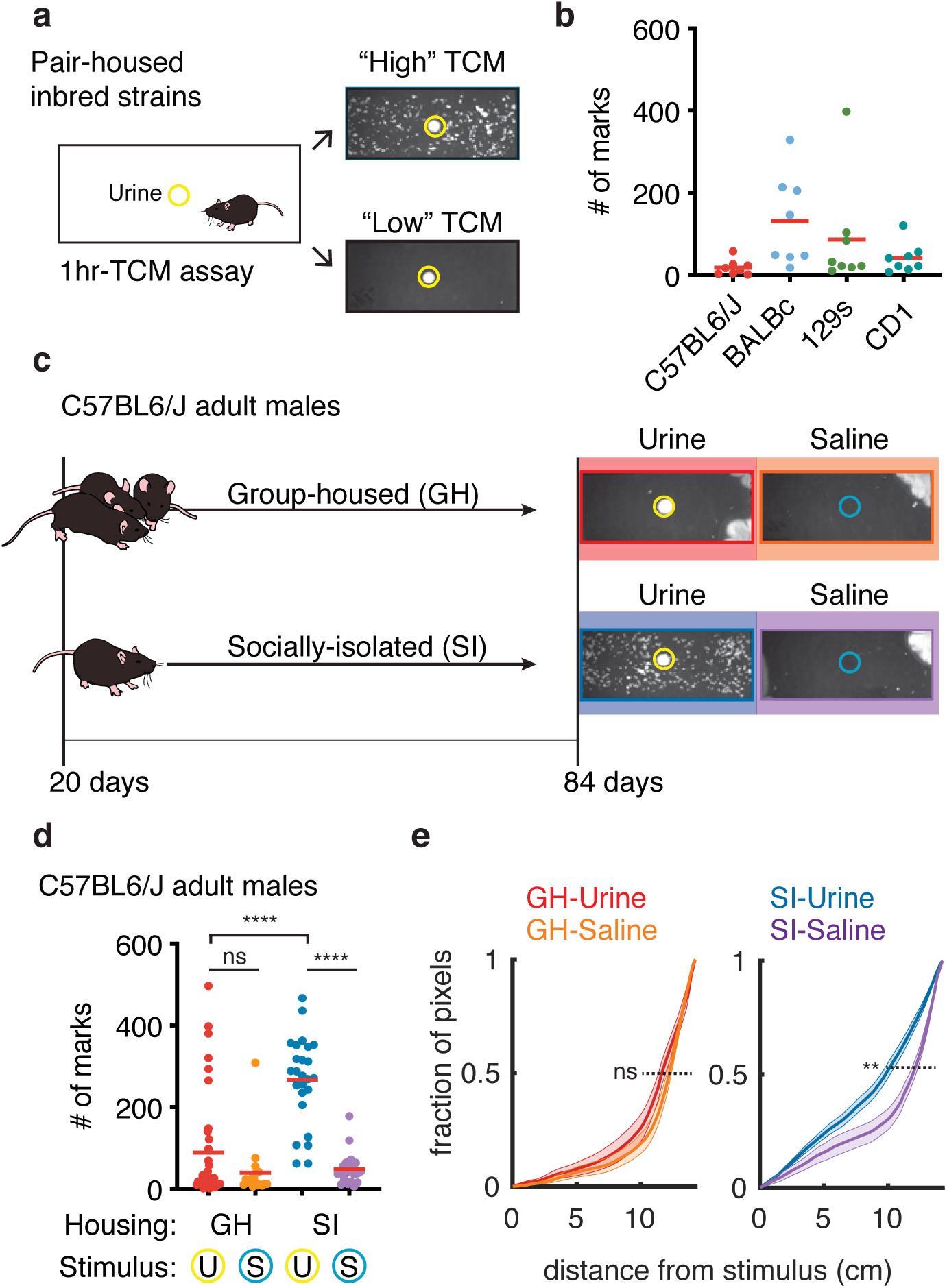
Social isolation reveals an innate form of context-dependent territory-covering micturition (TCM). a) Schematic of the assay used to analyze TCM of pair-housed mice from 4 different inbred strains. Each pair-housed mouse was individually placed in a filtered paper lined cage with urine stimulus (yellow circle) in the center. Representative “High” and “Low” TCM patterns revealed after 1 hour in the arena are displayed on the right. b) Total number of urine marks deposited by mice of each strain (n=8 mice per strain, red bar denotes the mean number of marks). c) Left: housing schematics for group-housed (GH) and socially-isolated (SI) animals. Right: Representative TCM patterns revealed 1hr after GH or SI animals were placed in a test chamber with saline (blue circle) or non-self urine stimulus (yellow circle). d) Total number of urine marks for each experimental condition (U=Urine exposed, S=saline exposed; n=36 mice for GH Urine, n=18 mice for GH Saline, n=25 mice for SI Urine, n=23 mice for SI Saline. ****p<0.0001; ns=not significant; two-tailed Mann-Whitney *U* test). e) Cumulative probability distribution of distances of urine-marked pixels from the stimulus center for the 4 contexts with mean (solid lines) and SEM (shaded areas). **p<0.01; ns=not significant; simulation by bootstrap analysis).

To assess the effect of social history on micturition behavior in male mice, we placed group-housed (GH) C57BL6/J males in cages with either a urine stimulus or a saline control. In both contexts, most C57BL6/J males either did not deposit urine or did so in several large spots near the corner of the cage (Fig. 1c). This pattern of urination has been previously associated with subordinate males and with females. A few male mice (6 out of 36 mice) displayed micturition behavior in which they covered the territory in the urine environment but not when exposed to saline (GH-Urine: 89 ± 22 marks; GH-Saline: 39±16 marks; n=36 and 18 mice; p=0.3 Mann-Whitney test, Fig. 1d).

We hypothesized that continuous social interactions in the group actively suppresses context-dependent TCM. The variability in individual experience and the changes they induce in each animal (e.g., stress and hormonal factors) may hinder the ability to robustly observe TCM and reveal the circuitry distinguishing social and sensory-motor components of the behavior. To test the hypothesis that TCM is altered by social experience, we socially isolated C57BL6/J males after weaning. Once the animals reached adulthood (>p84) (Fig. 1c), we monitored TCM as above in the presence of saline or urine. In contrast to GH animals, the majority of socially-isolated (SI) males produced extensive TCM in the presence of urine but not saline (SI-Urine: 266 ± 21 marks; SI-saline: 48 ± 8 marks; n=25 and 23 mice; p<0.0001 Mann-Whitney test; Fig. 1d). We observed the same pattern in the BALBc strain, but not in 129s and CD1 strains (Supplementary Fig. 1b). Thus, social isolation reveals robust context-dependent TCM in C57BL6/J and BALBc males.

To understand if the spatial distribution of urine marks is different when mice display week or robust TCM, we calculated the cumulative probability distribution (CDF) of the distances (range 0-15 cm) of urine marks from the center of the arena (defined as 0 cm and where the urine or saline stimulus was deposited) (Fig. 1e). Whereas all GH mice and the SI mice exposed to saline deposit urine far from the center of the cage (50% of CDF at GH-Urine 11.6±0.05 cm, GH-Saline: 12.2±0.05 cm, SI-Saline: 11.9±0.03 cm) the SI mice exposed to urine placed urine closer to the center (9.8 ± 0.00 cm. p=0.004 for SI-Urine and SI-Saline, p=0.1 for GH-Urine vs SI-Saline, test by bootstrap analysis).

Taken together, these results demonstrate that social isolation in C57BL6/J mice induces TCM. Therefore, this behavior paradigm can be used to isolate how sensory environment modulates the bladder output in TCM, bypassing its modulation by social interactions.

### Quantitative comparison of brain-wide inputs to *Crh+* PMC neurons

We previously identified a cluster of *Crh+* neurons in the PMC whose activity was sufficient to induce bladder contraction and urine release (Hou et al., 2016). We hypothesized that the differences in TCM behavior in GH and SI males might result from changes in the projections to *Crh+* PMC neurons. Differences in the circuitry between GH and SI animals could reveal networks upstream of the PMC that modulate bladder output in context-dependent TCM. We used whole-brain mapping of neurons labeled with retrograde trans-synaptic rabies virus to quantitatively compare the distribution of cells putatively presynaptic to *Crh+* PMC neurons in GH and SI C57BL6/J male mice (Fig. 2a). We performed similar analysis in C57BL6/J group-housed female mice to investigate potentially sexually divergent presynaptic areas that might contribute to this a male-specific behavior (Kimura and Hagiwara, 1985; Rosell and Thomsen, 2006).

**Figure 2.**
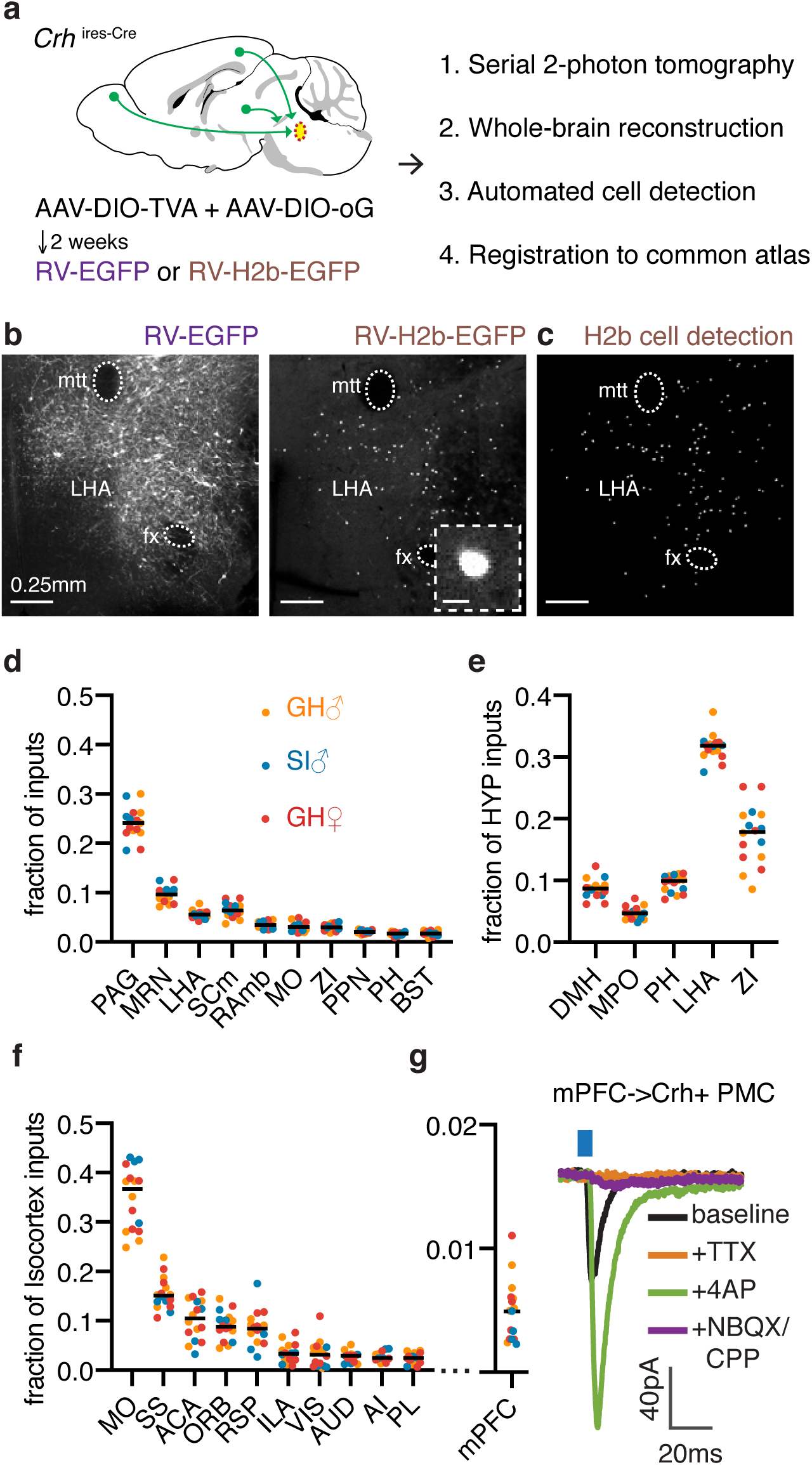
Stable brain-wide connectivity of the circuitry upstream of PMC across housing conditions and sex. a) Schematic of rabies-based monosynaptic retrograde trans-synaptic tracing from *Crh+* PMC neurons. AAV and RV injected brains were imaged with serial 2-photon tomography, reconstructed in 3D, and registered to a reference atlas for automated analysis. b) Representative coronal sections of cell-filing RV-EGFP labeled cells (left) and nuclear localized RV-H2b-EGFP labeled cell (right, inset with zoomed view of one RV labeled cell, scale bar = 10µm) in lateral hypothalamic area (LHA; mtt= mammillothalamic tract, fx=fornix). c) Output of the cell-detection algorithm from the same RV-H2b-EGFP hypothalamic section as in B. d) Top 10 putative inputs to *Crh+* PMC neurons, presented as the distribution of candidate neurons in GH-Male, SI-Male, and GH-Females (n=6 mice for GH-male, n=4 mice for SI-male, n=6 mice for GH-Female). Differences among the groups were not statistically significant (p>0.05 for all regions, Wilcoxon Signed Rank Test, adjusted for multiple comparisons with the Benjamini, Kreiger, and Yekutieli false discovery rate approach (FDR=0.05)). e) The distribution of top 5 hypothalamic inputs shown as the fraction of the total labeled cells within the hypothalamus. f) The distribution of top 10 cortical inputs shown as fraction of the total labeled cells within the isocortex. g) Monosynaptic glutamatergic connections of mPFC -> *Crh+* PMC (35/60 neurons). For acronyms of brain regions, refer to Supplementary Table 1

To achieve an unbiased quantification of the distributions of putative presynaptic connectivity to *Crh+* PMC neurons, we utilized trans-synaptic rabies tracing from genetically identified cell types with automated image acquisition and analysis (Ragan et al., 2012; Wickersham et al., 2006; 2007). Rabies viruses (RV) that express cell-filling fluorophores, such as GFP, highlight the full neuronal morphology including axons and dendrites (Fig. 2b). Although this, in theory, can provide more information about the GFP-expressing neuron, in practice the fluorescence from overlapping processes in the neuropil greatly complicates analysis. To allow simple image analysis to find all the RV-labeled cell bodies, we generated a rabies virus expressing nuclear-localized GFP (RV-H2b-EGFP). Tight nuclear localization of RV-H2b-EGFP allowed precise quantification of rabies-infected neurons, permitting an unbiased whole-brain estimate of putative inputs to *Crh+* PMC neurons across different housing conditions and sex (Fig. 2c). In contrast to cell-filling RV-EGFP, RV-H2b-EGFP yielded robust cell-detection due to the nucleus-restricted fluorescence (Mandelbaum et al., 2019).

Cre-dependent AAVs encoding TVA-mCherrry and rabies glycoprotein (RVG) were injected into the PMC of *Crh^-^*^ires-Cre^ mice to render Crh+ neurons sensitive to infection with EnvA pseudotyped RV and to complement the G-deleted RV to move trans-synaptically following infection (Kim et al., 2016; Wall et al., 2010). After 2 weeks, we injected RV-H2b-EGFP into the same coordinates (Fig. 2a). As control experiments for the cell-type specificity, injection of EnvA pseudotyped RV without prior AAV-DIO-TVA injection showed no rabies infection, whereas omitting AAV-DIO-RG resulted in robust starter cell labeling but no trans-synaptic labeling of putative inputs (Supplementary Fig. 2a-b).

The distribution of H2b-EGFP labeled neurons in the 3D brain volume was subsequently imaged through serial two-photon tomography (Ragan et al., 2012), reconstructed, and aligned to the Allen Brain Atlas (ABA). The fraction of total RV-H2b-EGFP cells was calculated in each ABA-defined region to quantify the distribution of RV+ labeled inputs (Supplementary Table 2). In the three groups with divergent TCM (n=6 GH males, n=4 SI males, n=6 GH females), the largest fraction of input cells originated in the periaqueductal grey (Fig. 2d; PAG, 24.2±0.6 % of total inputs), followed by midbrain reticular nucleus (MRN, 9.8±0.6% of total inputs).

We utilized the quantitative whole-brain rabies mapping dataset as a discovery tool to identify previously unknown components of the micturition network. Querying the Allen Brain Atlas Injection Map, we discovered that *Crh+* PMC neurons also send projections to ventrolateral part of PAG where putative inputs to *Crh+* PMC reside (Supplementary Fig. 2c). PAG receives ascending sacral afferent inputs (Fowler et al., 2008; Groat and Wickens, 2013) as well as projections from hypothalamic nuclei. Thus, the PAG-PMC loop might serve as an additional mechanism to control voluntary micturition.

Hypothalamic inputs accounted for 18% of total forebrain inputs to *Crh+* PMC neurons (Supplementary Fig. 2d), with the lateral hypothalamus (LHA)-zona incerta (ZI) complex providing about 50% of these (Fig. 2e; LHA: 31.5±0.5% of hypothalamic inputs, ZI: 17.4±1.1% of hypothalamic inputs). By comparison, the medial preoptic area (MPO), a hypothalamic nucleus that modulates TCM through the PMC (Hou et al., 2016), accounted for about 0.8% of total forebrain inputs and about 5% of hypothalamic inputs.

Other areas identified as putatively presynaptic to PMC *Crh+* neurons were the motor cortex (MO), which conveys top-down urination initiation signals to the PMC (Yao et al., 2018), and the medial prefrontal cortex (mPFC, ILA+PL), a region involved in the regulation of social dominance (Wang et al., 2011; Zhou et al., 2017). MO had the highest number of RV-labeled neurons among cortical areas (3.1±0.2% of total RV+ inputs, 6^th^ overall). Within cortex, MO provided 35% of all putative cortical inputs serving as the largest source of cortical input while the mPFC sent considerably less (Fig. 2f; <1% of total inputs; 5.5±0.5% of total cortical inputs, see Supplementary Table 2 for full dataset). Additionally, within the PMC extensive labeling of non-*Crh* neurons was observed, indicating potential intra-PMC connectivity (Supplementary Fig. 2e).

Subsets of identified putative inputs were tested for functional connectivity through *in vitro* electrophysiology. We injected Cre-independent ChR2 into candidate upstream regions of *Crh*^ires-Cre^*;Rosa26*^lsl-tdTomato^ animals in which *Crh*-expressing neurons are marked by tdTomato red fluorescence. Using whole-cell voltage-clamp recordings, we analyzed light-evoked postsynaptic currents in *Crh+* PMC neurons (Petreanu et al., 2007). We observed a clear monosynaptic glutamatergic current from mPFC (<1% of inputs), highlighting the sensitivity and accuracy of the RV-based trans-synaptic tracing (Fig. 2g, Supplementary Fig. 2f-i; 35/60 *Crh+* neurons were connected).

To test the hypothesis that circuit wiring differences contribute to the differences in TCM behavior, we compared the distribution of putative inputs to PMC *Crh+* neurons in GH males, SI males, and GH females. The brain-wide distributions of RV+ neurons were similar across all three conditions (p>0.05, Wilcoxon Signed Rank Test, adjusted for multiple comparisons with the Benjamini, Kreiger, and Yekutieli false discovery rate approach (FDR=0.05); Fig 2.d-f), suggesting that, despite the behavioral divergence, GH and SI males as well as GH females largely share a similar network upstream to the PMC.

### Social isolation changes brain-wide activity patterns

Although retrograde trans-synaptic rabies has been proposed to depend on both anatomy (i.e. the existence of a synapse) and functional properties (i.e. the strength of the synapse) (Beier et al., 2017), there are likely changes in synaptic properties of neurons that project to PMC that are not revealed by the distribution of RV-labeling. Furthermore, alterations in the intrinsic properties of Crh+ neurons might contribute to changes in activity of the PMC and micturition behavior. Therefore, we prepared acute brain slices from adult mice and compared intrinsic electrophysiological properties of *Crh+* PMC neurons in across social experience (Supplementary Fig. 3a). Current clamp recordings revealed only a moderate increase in the number of action potentials evoked by current injections in SI males, which results from decreased action potential threshold (Fig. 3b-c). No changes were observed in resting membrane potential, sag potentials, and membrane capacitance between the two groups (Supplementary Fig. 3b).

**Figure 3.**
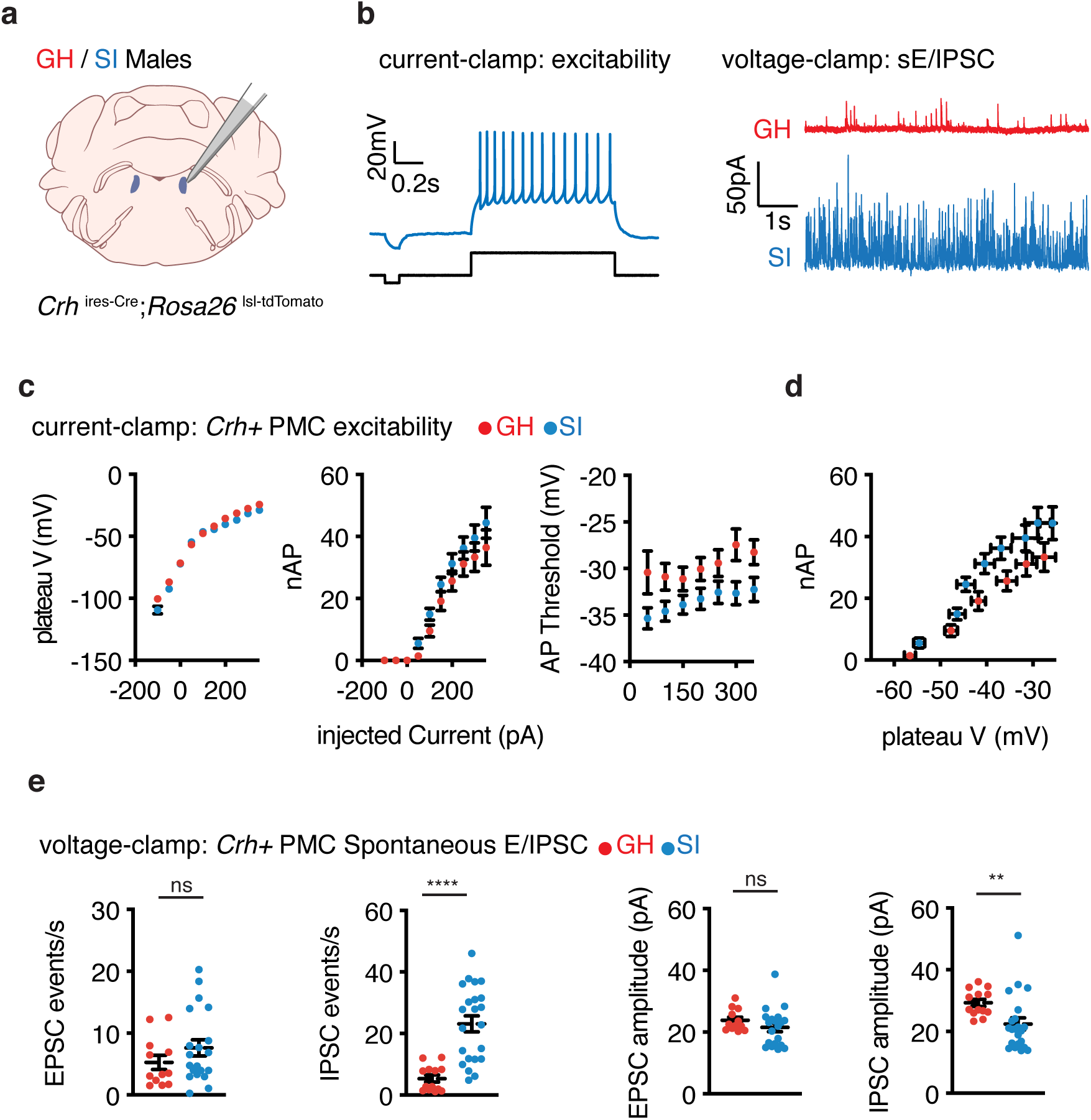
Social isolation induces synaptic changes at *Crh+* PMC neurons. a) Current-clamp and voltage-clamp recordings from *Crh+* PMC neurons used to assess intrinsic excitability and synaptic changes, respectively. b) Example traces of current clamp recording and voltage clamp recording. (Left): Example trace of current-clamp recording (top) with the current injected (bottom). (Right): Example traces of inhibitory post-synaptic currents (IPSC) in GH and SI males. c) Plateau potential, number of action potentials, and first action potential threshold potential as a function of injected (current amplitude ranging from −200 to 400pA). Analysis method is described in Supplementary Fig. 4. d) Comparison between the numbers of action potential and plateau potentials in *Crh+* PMC neurons between the GH (red) and SI (blue) animals (n=5 animals 35 cells from GH males, 4 animals and 29 cells from SI males). e) Frequencies and amplitudes of excitatory post-synaptic currents (EPSC) and IPSCs in Crh+ PMC neurons of GH and SI males (EPSC: n=4 animals 12 cells from GH males, n=5 animals 20 cells from SI males; IPSC: n=4 animals 13 cells from GH males, n=5 animals 22 cells from SI males; ns=not significant, ****P<0.0001, **P<0.01; two-tailed Mann-Whitney *U* test).

We also obtained whole-cell voltage-clamp recordings from *Crh+* PMC neurons in acute brain slices from adult GH and SI mice and compared spontaneous excitatory and inhibitory post-synaptic currents (Fig. 3d, sEPSCs and sIPSCs). The frequencies of both postsynaptic currents were higher in SI males, especially the sIPSCs (Fig. 3e; GH sEPSC: 5.2±1.1 Hz; SI sEPSC: 7.6±1.3 Hz, p=0.2; GH sIPSC: 5.5±1.2 Hz, SI sIPSC: 23.2±2.6 Hz, p<0.0001 Mann-Whitney test). Large changes in sIPSC rates suggest enhanced probability of release from inhibitory inputs to *Crh+* neurons or increased spontaneous activity of local inhibitory neurons in the PMC brain slice. Together with the rabies tracing data, this suggests that SI animals display synaptic changes upstream of *Crh+* PMC neurons without large-scale anatomical changes in the presynaptic network.

### Identification of upstream brain-wide micturition networks that are differentially regulated by context-dependent TCM

To identify brain regions with potentially differential activity that might contribute to the expression of TCM, we examined whole-brain patterns of c-fos protein expression following different experiences. As we had identified two features that regulate TMC – social experience and exposure to urine – we examined each of these separately.

First, group-housed and socially-isolated C57BL6/J males were exposed to urine in a behavioral arena as in Fig. 1. In this experiment, both sets of animals are exposed to the same sensory environment and only the previous social experience of mouse differs. One hour after exposure, the animals were euthanized and perfused and the brains cleared using iDisco+ (Renier et al., 2016). Subsequently, endogenous c-fos protein was immunolabeled and visualized using fluorophore-conjugated secondary antibody. Both far-red (c-fos) and green (autofluorescence) were detected via light-sheet imaging, followed by automated image-based identification of c-fos protein expressing cells and registration through the ClearMap pipeline (Fig. 4a-b; Full dataset in Supplementary Table 3).

**Figure 4.**
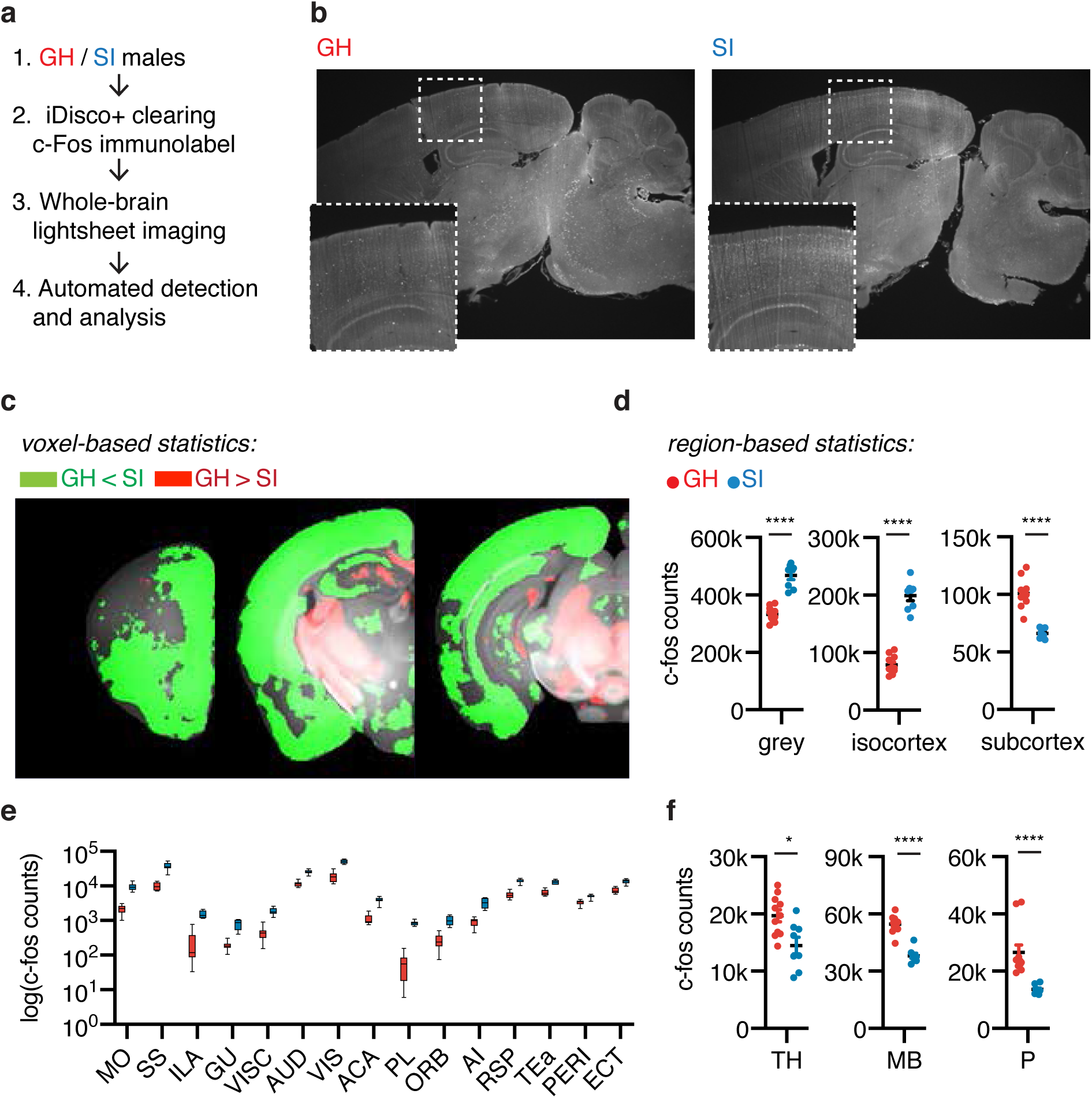
Social Isolation induces brain-wide activity changes. a) Whole-brain c-fos analysis workflow: brains from GH and SI males were cleared, immmunolabeled for c-fos, imaged, and analyzed. b) Representative sagittal view of c-fos immunolabled brains of group-housed and singly-housed animals. (bottom-left: inset zoomed view of cortex labeled with dotted boxes). c) Voxel-based statistics maps overlaid on Allen Brain Reference Atlas. Green indicate statistically significant (n=11 GH group, n=8 SI group; q<0.05 with FDR=0.05) voxels with higher c-fos counts in SI animals, and red indicate higher c-fos counts in GH animals. d) Quantification of c-fos cell counts of GH and SI males in grey, isocortex, and subcortex (n=11 GH group, n=8 SI group; Mann-Whitney test; ****P<0.0001). e) Quantification of c-fos cell counts of GH and SI males throughout isocortex presented in log scale. f) Quantification of c-fos cell counts of GH and SI groups in TH, MB, and P (Mann-Whitney test, *P<0.05, ****P<0.001). For full acronyms of brain regions, refer to Supplementary Table 1.

SI animals showed higher brain-wide c-fos+ cell counts (Fig. 4c-d; GH grey: 329846±7393, SI grey: 465235±14750, p<0.0001; Mann-Whitney test). Notably, the greater c-fos+ counts in socially-isolated animals was driven by increases in the isocortex (Fig. 4d; GH-isocortex: 73041±4649, SI-isocortex: 193785±8839, p<0.0001; Mann-Whitney test). Almost all sub-regions of the isocortex (Fig. 4e) exhibited a significant increase in the number of cells expressing c-fos, with the exception of subset of motor cortex previously associated with licking behaviors (ALM, Fig. 4c) (Chen et al., 2017). In contrast, ‘subcortical’ regions (Pons, Midbrain, Thalamus in ABA) displayed decreased number of c-fos+ cells compared to GH (Fig. 4d-f, Supplementary Fig. 4; GH-subcortex: 100854±3739, SI-subcortex: 66172±1592, p<0.00001; Mann-Whitney test). Thus, our data suggest that distributed and coordinated changes in activity across the brain, as opposed to in a single brain region, underlie the differences in behaviors of social-isolated and group-housed mice.

Second, we compared the whole-brain c-fos expression patterns of SI males exposed to urine or saline (Fig. 5a; n=4 mice each, Supplementary Fig. 5a). As expected, SI males exposed to urine marked the territory more robustly than SI males exposed to saline (Supplementary Fig. 5b). The distributions of endogenous c-fos protein induced by these different contexts were compared using ROI- and voxel-based statistical tests, corrected for multiple comparisons with a false discovery rate of 0.05. Voxel-based analysis revealed significant activation in the PMC of the urine-exposed mice, confirming previous findings (Fig. 5b). We also observed correlated network structures across multi-region groups, suggesting experience-specific brain-wide patterns of activation (Supplementary Fig. 5c).

**Figure 5.**
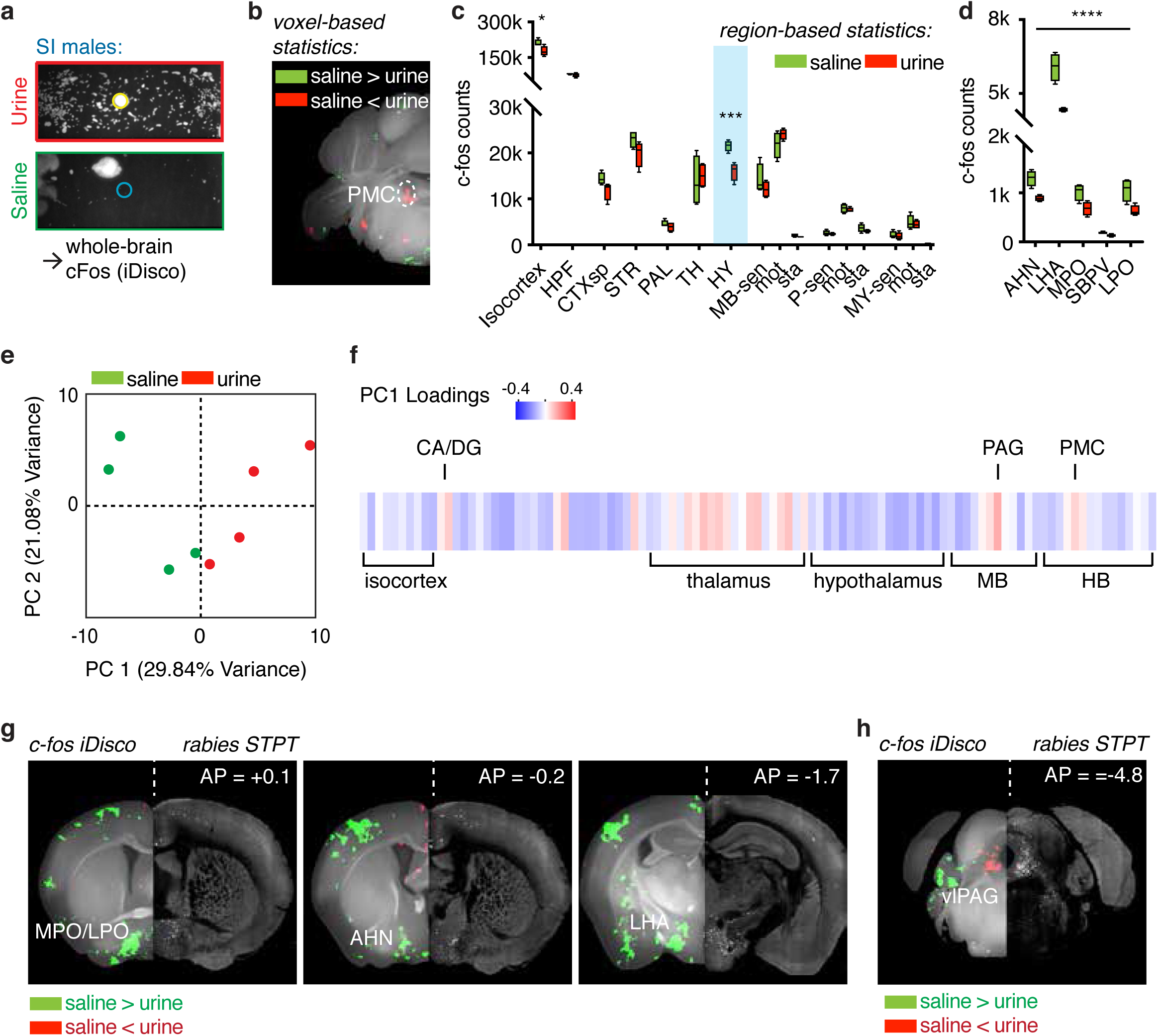
Different modes of TCM recruit differentially activated PMC upstream networks. a) Schematic for whole-brain c-fos analysis workflow: SI males were either exposed to urine or saline, which resulted in divergent TCM. 1 hour after the behavior, animals were perfused, the brains were cleared, immmunolabeled for c-fos, imaged, and analyzed (n=4 each for the urine and the saline conditions). b) Voxel-based statistics map reveals increase in the number of c-fos+ neurons in the PMC. Green indicate statistically significant (q<0.05, FDR=0.05) voxels with higher c-fos counts in animals exposed to saline, and red indicate higher c-fos counts in animals exposed to urine. c) c-fos induction differences in brain regions from saline and urine trials (*q<0.05, ***q<0.0001, FDR=0.05). d) Top 5 hypothalamic regions that displayed the most differences sorted by q-values (all regions q<0.0005; refer to Supplementary Table 2.3 for full statistics). e) Individual mice projected in the first two principal component (PC) space of whole-brain c-fos+ cell densities. f) Brain-wide c-fos patterns in response to urine, represented as individual regions’ loadings for the PC1. For acronyms of brain regions, refer to Supplementary Table 1. g) Hypothalamic regions with differential c-fos induction that are capable to project to the PMC: voxel-based statistics map from c-fos (left) and corresponding RV STPT images (right) of putative inputs to *Crh+* PMC in the same coronal slice. h) Ventrolateral segment of PAG, the major input to the PMC, displays differential c-fos activation.

Region-based analysis showed a large decrease in c-fos expressing cells in the hypothalamus in animals exposed to urine (Fig. 5c; HY; q<0.001 FDR=0.05). Many areas in the hypothalamus were differentially regulated, including regions previously implicated in socially-motivated behaviors such as lateral and medial preoptic areas (LPO and MPO) (Fig. 5d, Supplementary Fig. 7). Lateral hypothalamus (LHA), a major hypothalamic input to the PMC (Fig. 2e), displayed a significant decrease in c-fos labeled cells in urine exposed groups (q<10^-11^). Voxel-based analysis yielded identical results, with spatial patterns restricted to specific sub-hypothalamic nuclei. No differences were observed in ventromedial hypothalamus (VMH) (Supplementary Table 4), an area previously implicated in male aggression (Lin et al., 2011). Outside the hypothalamus, we observed that midbrain motor regions including subregions of the superior colliculus motor (SCm) and PAG were selectively activated in urine-exposed groups.

To investigate whole-brain network changes beyond single-region activation patterns, we applied principal component analysis (PCA) on the c-fos+ cell counts in urine and saline groups (DeNardo et al., 2019; Ye et al., 2016). Each mouse in the principal component space defined by the first two principal components (PCs) revealed that the urine and saline exposed groups segregated along PC1 (Fig. 5e). Examining the PC loadings across all the brain areas allowed us to visualize the brain-wide network pattern that separates the two groups (Fig. 5f). In general, the first PC was generated by negative weightings of cell counts in hypothalamus and positive ones in thalamus and midbrain. This indicates that the main differences between urine and saline exposed mice are explained by increase in c-fos labeling in thalamus and midbrain and decreases in hypothalamus.

Interestingly, hippocampal regions (dentate gyrus, DG; Ammon’s horn, CA) were recruited during in the urine exposure, suggesting potential spatial learning of the urine context (Fig. 5f). To test the hypothesis, two groups of male mice were prepared: one group was exposed to urine and displayed robust TCM, and the other group was exposed to saline and displayed low TCM. The next day, the two groups of mice were re-introduced to the same arena with no urine. In contrast to the mice exposed only to saline, mice previously exposed to urine continued to display robust TCM without re-exposure to the urine stimulus (Supplementary Fig. 5e-i). Since the TCM analysis of all mice was carried out in the same box, the differences in TCM cannot be explained by contaminating odors or other arena-specific cues.

To identify networks presynaptic to PMC that are potentially capable of modulating TCM directly, we computationally combined the rabies-tracing anatomical data with the c-fos iDisco immunostaining data. A side-by-side comparison of c-fos voxel maps and rabies cells revealed a striking spatial overlap (Fig. 5g-h). To obtain whole-brain comparison of structure and function of the presynaptic micturition network, we generated voxelated maps of rabies inputs on the same coronal atlas based on ABA (Supplementary Fig. 6). Notably, c-fos activation was seen in a region that borders anterior hypothalamus (AHN) / bed nuclei of stria terminalis (BST) but that is not well-defined by ABA; however, once referenced to the spatial input map labeled by rabies, the spatial pattern match is clear (Fig. 5g and Supplementary Fig. 6).

In summary, through unbiased and automated structure and function screening, we identified a putative upstream PMC micturition network that modulates TCM in context-dependent manner.

### Identification of critical networks upstream of the PMC underlying context-dependent TCM

Our data suggest that LHA is a robust input to PMC implicated in social-isolation and urine-induced TMC behavior – LHA is the major hypothalamic gateway to the PMC and has fewer c-fos labeled cells in mice displaying TCM. *In situ* hybridization revealed that the rabies-labeled neurons in LHA largely express genes encoding for glutamate decarboxylases (including both *Gad1* and *Gad2,* 66%, n=337 cells from 3 mice, Fig 6a-c), suggesting that LHA sends mostly GABAergic input to *Crh+* PMC neurons, potentially inhibiting micturition. If LHA does inhibit PMC and TCM, then suppression of the inhibition from LHA should elicit TCM in a saline context, and activation of the inhibition from LHA should inhibit TCM in a urine context.

**Figure 6.**
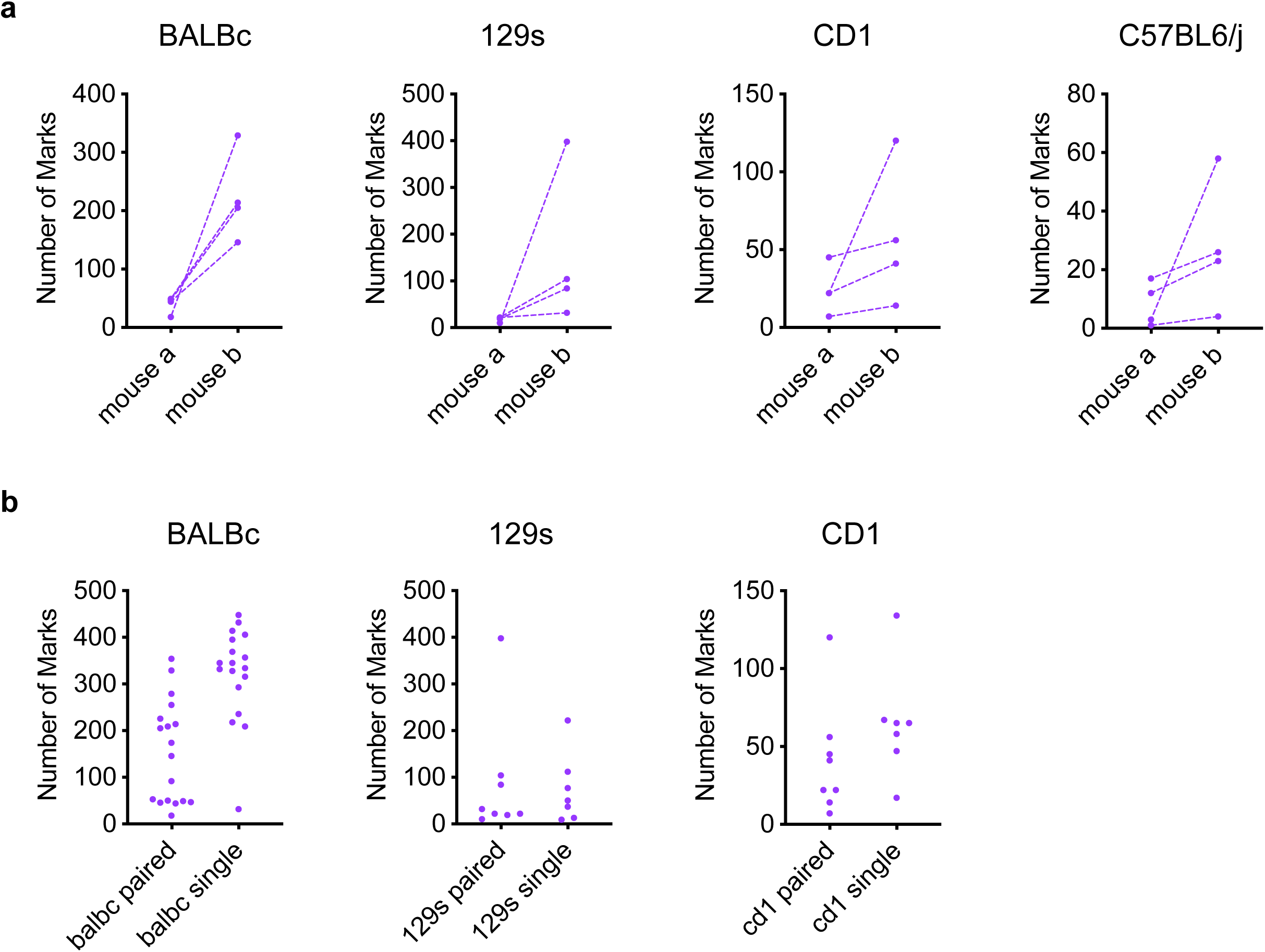
Bidirectional modulation of TCM via chemogenetic manipulation of the lateral hypothalamic area. a) Experimental design to interrogate neurotransmitter content of *Crh+* PMC projecting LHA neurons. Cell-type specific rabies tracing (described detail in Fig. 2) is performed, and sections including LHA were cryosectioned and cells were classified as GAD+ (*Gad 1 and Gad 2)* /RV (Rabies virus nucleoprotein) + or GAD-/RV+ through *in-situ* hybridization (ISH). b) Representative ISH image containing LHA. Arrowheads indicate *RV-N*+ cells. c) *in situ* hybridization of GAD and RV in LHA reveals that a majority of cells in LHA retrogradely labeled from *Crh+* PMC cells express GABA synthetic enzymes (n=3 mouse, 337 LHA cells). d-e) Experimental design: d) AAV encoding inhibitory DREADD hM4Di or excitatory DREADD hM3Dq was bilaterally injected in lateral hypothalamic area. e) 3 weeks after surgery, hM4Di mice were introduced to a test cage with no urine stimulus, and hM3Dq mice were introduced to a cage test with no saline stimulus. Mice were IP injected with either vehicle or CNO 25 minutes before the behavior. f) Left: representative expression pattern of DREADD targeted towards lateral hypothalamus, Right: representative image showing DREADD-expressing fibers in the pons. PMC is identified (oval) relative to the fourth ventricle (4v). g) Example TCM pattern from mice transduced with DREADD or mCherry control (left: vehicle injection, right: 5mg/kg CNO injection). h) Left: Number of urine marks deposited were increased in CNO trials compared to vehicle trials only in hM4Di transduced animals (n=6 hM4Di animals, n=4 mCherry controls, two-tailed Mann-Whitney test, **P<0.01, ns=not significant) Right: Number of urine marks deposited were decreased in CNO trials compared to vehicle-injected trials only in hM3Dq transduced animals (n=6 hM3Dq animals, n=5 mCherry controls, two-tailed Mann-Whitney test, **P<0.01, ns=not significant)

To test this hypothesis, we used the chemogenetic tools hM4Di and hM3Dq (Armbruster et al., 2007; Urban and Roth, 2015), which are engineered Gq and Gi, respectively, coupled receptors that are activated by the ligand clozapine-N-oxide (CNO) to either increase or reduce the activity of receptor-expressing neurons (AAV-hM4Di-mCherry or AAV-hM3Dq-mCherry, Figure 6d). hM4Di may also function by decreasing the probability of action potential-evoked neurotransmitter release from expressing neurons (Stachniak et al., 2014).

To test the first prediction that inactivation of LHA would elicit TCM in a saline context, we placed SI male mice in clean cages lined with filter paper and a saline stimulus. 30 minutes before the behavioral analysis, the animals were IP injected with either CNO (5mg/kg) or vehicle (Fig. 6e). Mice expressing hM4Di in lateral hypothalamus displayed increased TCM following CNO injection compared to after saline injection (Fig. 6g-h; n=5 hM4Di mice, hM4Di Vehicle: 56 ± 33 marks, hM4Di CNO: 173 ± 16 marks, p=0.043; Mann-Whitney test). mCherry-expressing control animals showed similar TCM on CNO and saline injection days (n=4 mCherry mice; mCherry Vehicle: 26 ± 4 marks, mCherry CNO: 12 ± 2 marks, p=0.93; Mann-Whitney test).

To test the second prediction that activation of LHA should inhibit TCM in a urine exposure context, we placed the animals in clean cages lined with filter paper with urine stimulus. 30 minutes before the behavioral analysis, the animals were IP injected with either CNO (5mg/kg) or vehicle. mCherry-expressing control animals displayed robust TCM on both saline and CNO injected days (Figure 6g-h; n=5 mCherry mice; mCherry Vehicle: 231 ± 28 marks, mCherry CNO: 429 ± 13 marks, p=0.86; Mann-Whitney test). In contrast, mice expressing hM3Dq in lateral hypothalamus dramatically decreased TCM in CNO-injected days, despite the urine stimuli (Figure 6g-h; n=6 hM3Dq mice; hM3Dq Vehicle: 259 ± 54 marks, hM3Dq CNO: 37 ± 7 marks, p=0.002; Mann-Whitney test).

Together, these results demonstrate that the activation and inactivation of LHA bidirectionally modulates TCM. Thus, LHA is one of critical nodes in the PMC micturition network that were identified with a series of brain-wide screening experiments.

## Discussion

Animals select specific motor actions based on sensory information about their current environment, memories of past experiences, and internal states. Most non-reflex behaviors that are influenced by these factors are complex. This precludes directly linking neural activity to specific muscle action, and obfuscates the flow and transformation of information from sensory organs to motor effectors. Studying the neural circuits underlying TCM is attractive for understanding decision-making because: 1) the behavior is simple (the bladder has two modes—retention and voiding), and culminates in an experimentally observable decision; 2) the decision is expressed with only three muscles whose activities are coordinated; and 3) the behavior is controlled by a sole effector nucleus, the pontine micturition center (PMC), onto which pathways that contain relevant information must converge.

### Social-hierarchy dependent TCM in the laboratory

Rodent social-hierarchy dependent TCM was first studied in outbred wild strains (Desjardins et al., 1973). Many follow-up studies have examined micturition in variety of inbred/outbred strains (Arakawa et al., 2008; Hou et al., 2016; Hurst, 1990; Kaur et al., 2014; Keller et al., 2018; Wu et al., 2009; Yao et al., 2018). Despite the advances that these studies brought, this experimental divergence prevented synthesizing behavioral findings across laboratories. Here, we systematically characterized TCM in different strains and housing conditions in order to understand the variability in micturition behavior and, hopefully, standardize approaches across laboratories.

Group-housed males live in a complex social structure—individual mice continuously interact with casemates, competing for resources (water/chow) and establishing a tiered social hierarchy. In addition, each social interaction is capable of inducing transcriptional (Williamson et al., 2018), electrophysiological (Golden et al., 2016; Zhou et al., 2017), and neuroendocrine (Lee et al., 2019; Williamson et al., 2017a; 2017b; 2019) changes in the brain. One example of a behavior influenced by social structure is TCM, for which is it well-established that social experience and the resulting social hierarchy play a significant modulatory role (Desjardins et al., 1973; Hou et al., 2016).

We hypothesize that the lack of robustness in TCM in group-housed C57bl6 is due to the complexities of continuous social interaction, and therefore, social isolation bypasses the challenges of experimentally monitoring and controlling social dominance. We propose that this provides a platform to understand the neural processes that underlie the sensorimotor transformation during TCM.

### Combined structure-function elucidation of TCM controlling brain centers

We find that that the combined analysis of circuit architecture with trans-synaptic labeling and identification of behavior-specific brain activation can reveal functionally relevant presynaptic micturition networks underlying TCM. Structural analysis alone failed to reveal circuit differences that explain varying TCM. The distribution of putative presynaptic inputs to *Crh*+ PMC neurons in group-housed males, singly-housed males, and females —groups with divergent TCM— differed only minimally. Previously, a nearly identical approach has been utilized to uncover connectivity differences of hypothalamic circuitry in males and females (Kohl et al., 2018) and across subregions of small thalamic nuclei (Mandelbaum et al., 2019), suggesting that we are not limited by our detection technology. Although we cannot account for unknown differences that are beyond the resolution of our methods—such as subtle synaptic plasticity or morphological changes—it is evident that the three groups share largely comparable structural connectivity of the upstream PMC micturition network.

Instead, we discovered changes in brain-wide activity between TCM conditions. These findings are consistent with studies in head-restrained mice that revealed distributed brain-wide representations of behavioral variables (Allen et al., 2017; Steinmetz et al., 2018; Stringer et al., 2018) that are gated by behavioral states (Allen et al., 2019). In contrast, the whole-brain c-fos induction pattern between different sensory and behavioral contexts within an environmental condition (singly-housed urine vs. saline) revealed relatively discretely affected networks throughout the whole-brain. Combined with multi-modal analysis of brain-wide rabies-tracing data, our work identified critical nodes in the network for the behavior conveying significant task-specific information that guide context- and state-dependent TCM via the PMC, including the lateral hypothalamic area.

Brain-wide structural and functional analyses suggest a simple rule behind neural control of TCM: social-environmental context is represented by coordinated global activity changes while TCM-relevant behavioral and sensory context are represented by discrete networks. This contrasts with the alternative hypothesis that individual nodes in the presynaptic PMC micturition network encode distinct state or sensory information. Practically, our data illustrate the need to investigate coordinated circuits throughout the whole-brain rather than studying one single specialized brain regions. Leveraging new technologies to record single-neuronal level activity throughout the large brain volumes (Jun et al., 2017; Sofroniew et al., 2016) together with the extensive connectivity atlas dataset (i.e. Allen Brain Connectivity Mapping) can shed light as to how whole-brain connectivity and activity dynamics can give rise to globally coordinated control of context-dependent TCM.

### PMC as an integrating switch of converging presynaptic micturition network

Our findings reaffirm the hypothesis that the PMC functions as an integration center of various pro- and anti-micturition cues converging onto the PMC that convey the environmental context and the state signals. Previously we reported that silencing of the medial preoptic area, one of the upstream micturition networks of the PMC, can mask social-rank dependent TCM. Here, we report bidirectional modulation of TCM through manipulation of the lateral hypothalamic activity. One thing to note is that the spatial pattern of TCM is dramatically different in the two experiments: inhibition of LHA led to an increase in TCM that extensively covers the territory whereas inhibition of MPO led to big circular urine spots (Hou et al., 2016). This suggests that the two different nodes of the presynaptic PMC network—LHA and MPO—encode and transfer different information to the PMC and differentially impact patterns of activity in PMC. This hypothesis is further supported by recent findings that motor cortical input to the PMC is critically involved in ‘micturition initiation’ (Yao et al., 2018). The activity of motor cortical input to the PMC increases its activity only during the initiation of each micturition bout, whereas the PMC sustains the increase in the activity throughout the micturition event (Yao et al., 2019).

This inspires future studies on how the intra-nuclear connectivity of PMC (Supplementary Fig. 2e) processes disparate inputs and translates each to micturition events. Multiple excitatory cell-types within the PMC may separately modulate the sphincter and the bladder wall, as reported by the studies of non-*Crh* expressing glutamatergic populations in the PMC (Keller et al., 2018). Recurrent connectivity within those glutamatergic cell types can confer rapidly coordinated and prolonged contraction of bladder wall and relaxation of urinary sphincter. Activity of local interneurons may mediate the switch between different modes of the PMC in different behaviors. These interneurons might be a target for neuromodulation that encodes information about behavioral state and modulates TCM in a context- and state-dependent manner. This hypothesis can be tested, refuted, and updated by a detailed survey of heterogeneity in the PMC, their local connectivity, and the network activity during micturition and the TCM.

## Materials and Methods

Further information and requests for reagents may be directed to, and will be fulfilled by, Bernardo L. Sabatini

### Mice

We used the following mouse lines in the study: Wild-type C57BL6J (Jackson Laboratory #000664). Wild-type BALBc (Jackson Laboratory #000651) mice. Wild-type 129s (Jackson Laboratory #002448) mice. Wild-type CD1 (Charles River #022) mice. Knock-in mice with an internal ribosome entry site (ires)-linked Cre recombinase gene downstream of the *Crh* locus (*Crh*^ires-Cre^, Jackson Laboratory # 012704) (Madisen et al., 2009; Taniguchi et al., 2011). Cre-dependent tdTomato (*ROSA*^lsl-tdTomato^, Jackson Labs # 007914) reporter mice (Madisen et al., 2009). All mice used in this study were between 2-5 months in age. Animals were kept on a 12:12 light/dark cycle or a reversed cycle under standard housing conditions. All experimental manipulations were performed in accordance with protocols approved by the Harvard Standing Committee on Animal Care, following guidelines described in the National Institutes of Health Guide for the Care and Use of Laboratory Animals. All mice brain coordinates in this study are given with respect to Bregma; anterior–posterior (A/P), medial–lateral (M/L), and dorsal–ventral (D/V).

### Virus Preparation

Recombinant adeno-associated viruses (AAV) of serotype 1, 5 and 9 encoding downstream gene under the control of CBA, Ef1a, or hSyn promoter was used throughout the study. Details of the viruses including specific promoters and serotypes are mentioned in individual method sections. All the AAVs were used at the concentration of 10^12^ GC/mL and was purchased from commercial vector cores (UNC, Penn, and AddGene). EnvA pseudotyped rabies viruses (RV-EGFP, RV-H2b-EGFP) were generated in-house as previously described (Mandelbaum et al., 2019; Wickersham et al., 2010).

### Stereotaxic intracranial injections and fiber optic implantation

All surgery was maintained in asceptic conditions. Mice were anesthetized with 2-3% isoflurane and placed in a stereotactic frame (David Kopf Instruments). Skulls were exposed and small holes were drilled into the skull. Viruses were injected (100-200nL total volume) at a rate of 100nL min^-1^ via glass pipettes with tip size of approximately 40um. Mice were given pre- and post-operative oral carprofen (MediGel CPF, 5mg/kg/day) as an analgesic, and monitored daily for at least 4 days post-surgery. All coordinates that were used in this study were relative to Bregma (in mm) and were: for mPFC: 2.1 A/P, 0.35 M/L, 2.0 D/V; LHA: −1.5 A/P, 1.1 M/L, 5.2 D/V; PMC: −5.3 A/P, 0.68 M/L, 3.5 D/V; LHA: −1.5 A/P, 1.1 M/L, 5.1 DV.

### Social isolation

Postnatal day 15-20 (p15-20) wild-type mice of 4 different strains were purchased from Jackson Labs and Charles River. At postnatal day 21, the mice were separated from the moms and housed in groups of 2-3 (pair-housed or group-housed), or in singles (singly-housed). The mice were housed in fully ventilated cages, and the handling was minimized. Individual cages were changed every two weeks.

### Territory-covering micturition (TCM) behavior

Adult (older than 70 days old) male mice were separated into individual fresh cages lined with filter paper (28 x 11 cm, Whatman #05-714-5) with an olfactory stimulus (50 µl male urine or saline) added to the center. Male urine is freshly collected and pooled from urine of 16-20 group housed BALBc males (for C57BL6/J, 129s, CD1 behavior) or C57BL6/J males (for BALBc males). After one hour in the arena, mice were removed, the distribution of urine spots on the paper was examined with fluorescence imaging as previously described (Hou et al., 2016). To quantify spatial distribution of urine marks throughout the cage, the urine mark image is thresholded and converted into a mask, which is used to calculate the distance distribution of all urine-covered pixels to stimulus center. Statistical significance was judged by simulation—for each housing condition (GH or SI) all the spatial CDF was put into one list, and randomly assigned to 2 groups. P value was calculated by how often one observes the differences out of 1000 re-runs.

For real-time tracking experiment (Supplementary Figure 5), adult wild-type BALBc male mice were habituated for 3 days, 20 minutes per day, in a custom-made arena with simultaneous real-time urine-deposit and positional tracking (Hou et al., 2016). After habituation, TCM was assayed by placing the mice in the test arena with 50µl of either male urine or saline in three days in a row. Urine marks were analyzed with fluorescence imaging as described above. Behavioral data was analyzed using a commercial software (Ethovision, Noldus).

### Whole-brain rabies tracing

TVA, a receptor of an avian virus envelope protein (EnvA), and rabies glycoprotein (RG) were introduced specifically in *Crh+* PMC neurons through two Cre-dependent viruses AAV9-CBA-DIO-TVA-mCherry and AAV9-CBA-DIO-oG) injected unilaterally (100nL of 1:1 mixture). After two weeks of expression, 200nL of RG-deleted (ΔRG) rabies virus pseudotyped with EnvA (RV-EGFP or RV-H2b-EGFP) was injected intracranially. RV injected animals were perfused transcardially with ice-cold PBS followed by 4% PFA, 7 days after RV injection. After a 24 hour post-fix in 4% PFA, brains were kept in 0.7% glycine solution for 48 hours. The brain was stored in PBS before embedding in 4% agarose in 0.05M PB, cross-linked in 0.2% sodium borohydrate solution, and imaged with high-speed multiphoton microscope with integrated vibratome sectioning (x-y resolution of 1 µm; z-step of 50 µm; TissueCyte 1000, TissueVision) as described before (Ragan et al., 2012). The raw image files were corrected for illumination, stitched in 2D, and aligned in 3D. EGFP-positive neurons automatically detected by a convolutional network trained to recognize cytoplasmic neuronal cell body labeling (Turaga et al., 2010) were visually validated, reconstructed in 3D, and their spatial information was registered by affine followed by B-spline transformation using the software Elastix (Klein et al., 2010) to a 3D reference brain based on the Allen Brain Atlas (Kim et al., 2015; Sunkin et al., 2013). The number of total input neurons in the brain was normalized by the total number of RV positive neurons. Parts of P-pons, H-hindbrain, and MOB-main olfactory bulb were damaged or missing during the dissection and excluded from the analysis.

### Electrophysiology

Acute slice electrophysiology experiments were done as previously described (Saunders et al., 2015; Wallace et al., 2017). 300µm slices were cut in ice-cold solution containing (in mM) 25 NaHCO3, 25 Glucose, 1.25 NaH2PO4, 7 MgCl2, 2.5 KCl, 0.5 CaCl2, 11.6 ascorbic acid, 3.1 pyruvic acid, 110 Choline chloride. Slices were transferred for 10 min to a holding chamber containing choline-based solution consisting of (in mM): 110 choline chloride, 25 NaHCO3, 2.5 KCl, 7 MgCl2, 0.5 CaCl2, 1.25 NaH2PO4, 25 glucose, 11.6 ascorbic acid, and 3.1 pyruvic acid before transferring to a second room temperature chamber with ACSF for at least 30 min. Recordings were performed at 32 °C in carbogen bubbled ACSF using Cs-based internals for voltage-clamp measurements (in mM: 135 CsMeSO_3_, 10 HEPES, 1 EGTA, 3.3 QX-314 (Cl^−^ salt), 4 Mg-ATP, 0.3 Na-GTP, 8 Na_2_-Phosphocreatine, pH 7.3 adjusted with CsOH; 295 mOsm·kg^−1^) and K-based internals for current-clamp measurements (in mM: 135 KMeSO_3_, 3 KCl, 10 HEPES, 1 EGTA, 0.1 CaCl_2_, 4 Mg-ATP, 0.3 Na-GTP, 8 Na_2_-Phosphocreatine, pH 7.3 adjusted with KOH; 295 mOsm·kg^−1^). For optogenetics experiments, 100nL mixture of AAV5-Ef1a-DIO-ChR2 and AAV1-hSyn-Cre was co-injected in the mPFC. 5ms duration light pulses from a 473 nm laser 5-10mW per mm^2^ measured at the sample plane were used.

### Electrophysiology analysis

All analysis regarding electrophysiological properties were performed using custom-written MATLAB script. In short, action potentials were identified from peaks crossing 0 mV. Action potential threshold was determined from the voltage at the time corresponding to the peak of the second derivative of the voltage and maximum dV/dT was measured from the peak of the first derivative. For ChR2-evoked EPSC analysis, resting membrane potential was measured as the median of potentials during periods of the sweep that had no current injection, and the peak post-synaptic current was measured manually. For spontaneous EPSC/IPSC analysis, candidate EPSC/IPSC amplitude was identified by findpeaks function in MATLAB and was manually verified in by an experimenter.

### Whole-brain c-fos staining and imaging

Single-housed adult C57BL6/J males and pair-housed adult C57BL6/J males (older than 70 days old) were separated into individual fresh cages lined with filter paper (28 x 11 cm, Whatman #05-714-5) with an olfactory stimulus (50 µl male urine or saline) added to the center. After one hour in the arena, mice were returned to the home cage. One hour later, the mice were deeply anesthetized with isoflurane and transcardially perfused with ice-cold saline followed by 4% PFA. After 24-hour post-fix in 4% PFA, brains were kept in 0.05M PB solution.

Within 3 weeks of perfusion, the left hemispheres of the brain samples were cut and were immunolabeled for c-fos and subsequently cleared using iDisco+ protocol (Certerra, NY; Renier et al., 2016). In short, the samples were pretreated with methanol, permeabilized with solution containing DMSO, blocked in donkey serum, and passively immunolabeled for c-fos (9F6 Rabbit mAb #2200, Cell Signaling Technology) and Alexa Fluorophore conjugated 647 secondary antibodies. Then, the samples were cleared with a combination of methanol and dichloromethane as previously described (Renier et al., 2016).

### Whole-brain c-fos analysis

Cleared samples were imaged in sagittal orientation (right lateral side up) on a light-sheet fluorescence microscope (Ultramicroscope II, LaVision Biotec) equipped with a sCMOS camera (Andor Neo) and a 4x/0.5 objective lens (MVPLAPO 4x) equipped with a 6-mm working distance dipping cap. The samples were scanned with a step-size of 3 micrometers using the continuous light-sheet scanning method with the included contrast blending algorithm for the 640 nm and 595 nm channels (20 acquisitions per plane), and without horizontal scanning for the 480-nm channel.

The activated c-fos+ neurons were automatically computationally identified and visualized in 3D. The datasets were warped in 3D by affine and B-spline transformation to an average Reference mouse brain generated from forty 8-week old C57BL/6 brains, as described previously (Kim et al., 2016a; Wall et al., 2010)

Statistical comparisons between different groups are run based on either ROIs or evenly spaced voxels. Voxels are overlapping 3D spheres with 100 µm diameter each and spaced 20 µm apart from each other. The cell counts at a given location, Y, are assumed to follow a negative binomial distribution whose mean is linearly related to one or more experimental conditions, X: E[Y]=α+βX. For example, when testing an experimental group versus a control group, the X is a single column showing the categorical classification of mouse sample to group id, i.e. 0 for the control group and 1 for the experimental group (O’Hara and Kotze, 2010; Venables et al., 2002). We found the maximum likelihood coefficients α and β through iterative reweighted least squares, obtaining estimates for sample standard deviations in the process, from which we obtained the significance of the β coefficient. A significant β means the group status is related to the cell count intensity at the specified location. The p-values give us the probability of obtaining a β coefficient as extreme as the one observed by chance assuming this null hypothesis is true. To account for multiple comparisons across all voxel/ROI locations, we thresholded the p-values and reported false discovery rates with the Benjamini-Hochberg procedure (FDR=0.05)

### *In situ* hybridization

For fluorescent *in situ* hybridization, animals were anesthetized with isoflurane before decapitation, and the brains were rapidly removed and frozen. 20µm PMC coronal sections of freezing media embedded brains (Tissue-Tek O.C.T) were prepared on cryostat (Leica, CM1950), and mounted on SuperFrost Plus glass slides (VWR) at 60µm intervals. Multiplexed fluorescent *in situ* hybridization was performed using the ACDBio RNAScope reagents and protocols as previously described (Hou et al., 2016). Single-plane images of the PMC were acquired via confocal microscope (Leica SP8).

### Chemogenetic Manipulation

hM3Dq (excitatory DREADD) or hM4Di (inhibitory DREADD) were non-Cre dependently introduced in bilateral LHA and nearby regions (AAV5-Ef1a-DO-hM3Dq, or AAV9-Ef1a-DIO-hM4Di mixed with AAV1-hSyn-Cre, 150nL per side) of BALBc wild-type single housed mice. For the control mice, AAV encoding mCherry was injected instead (AAV5-hSyn-mCherry). All behavioral experiments were conducted between 3-5wks post-injection. All mice were habituated for i.p injections for 2 days. Then the mice were subjected to behavioral testing: 30 minutes before the behavior, the mice underwent IP injection of CNO (5mg/kg) or volume-matched saline in the home cage. Then the mice were transferred to individual cages with either male urine (hM3Dq) or saline (hM4Di) in the center of the paper for 1 hour. The filter paper was then imaged and analyzed as described above.

After behavioral experiments, animals were perfused transcardially with ice-cold phosphate buffered saline (PBS) followed by 4% paraformaldehyde (PFA). After an overnight fix in 4% PFA, brains were equilibrated in 30% sucrose for at least 48 hours. Brain samples were then sliced in 50 µm using a frozen microtome (Leica) and every other section was mounted on Superfrost Plus (Fisher Scientific) slides. Slides were coverslipped with Prolong Antifade mounting media containing DAPI (Molecular Probes) and imaged with an Olympus VS120 slide-scanning microscope using 10x objective. Injection sites were manually confirmed based on the DREADD-mCherry expression

### Quantification and Statistical Analysis

Data points are stated and plotted as mean values ± SEM. p values are represented by symbols using the following code: * for 0.01<p<0.05, ** for 0.001<p<0.01, *** for 0.0001<p<0.001, and **** for p<0.0001. Exact p-values and statistical tests are stated in figure legends and methods. No *a priori* power analyses were done.

## Acknowledgement

We thank J. Saulnier, L. Worth, L. Chung, A. Philson, K. Robertson for their technical and administrative support; S. Knemeyer for help with the illustrations; L. Qi for help with the learning behavior; L. DeNardo for help with whole-brain c-fos analysis; C. Chen, C. Harvey, M. Andermann, D. Lin, L. Orefice, S. Liberles, T. Anthony, Y. Kim, and members of the Sabatini laboratory for helpful discussions and comments on the manuscript. Starting materials for generating RV are a generous gift from B.K. Lim (UC San Diego). Confocal images were acquired at the Harvard Neurobiology Imaging Facility (NINDS P30 Core Center Grant #NS072030). This work was supported by a Samsung Fellowship (M.H), a Lefler Predoctoral Fellowship (M.H.), a HMS Dept of Neurobiology Graduate Fellowship and a Stuart & Victoria Quan Fellowship (K.W.H.), and by the NIH (5R01DK114834-03 to B.L.S and P.O).

## Author Contributions

Conceptualization, M.H. and B.S.; Methodology: M.H., J.T., P.O., B.S.; Investigation: M.H., J.T., G.R., L.M., W.W., N.O., B.S.; Formal Analysis: M.H., J.T., L.M., P.O., B.S.; Resources: J.T., K.W.H., P.O.; Writing – Original Draft: M.H., B.S.; Writing – Review and Editing: All Authors; Supervision: B.S.; Funding Acquisition: B.S.

## Supplementary Figures and Tables

**Supplementary Figure 1.**
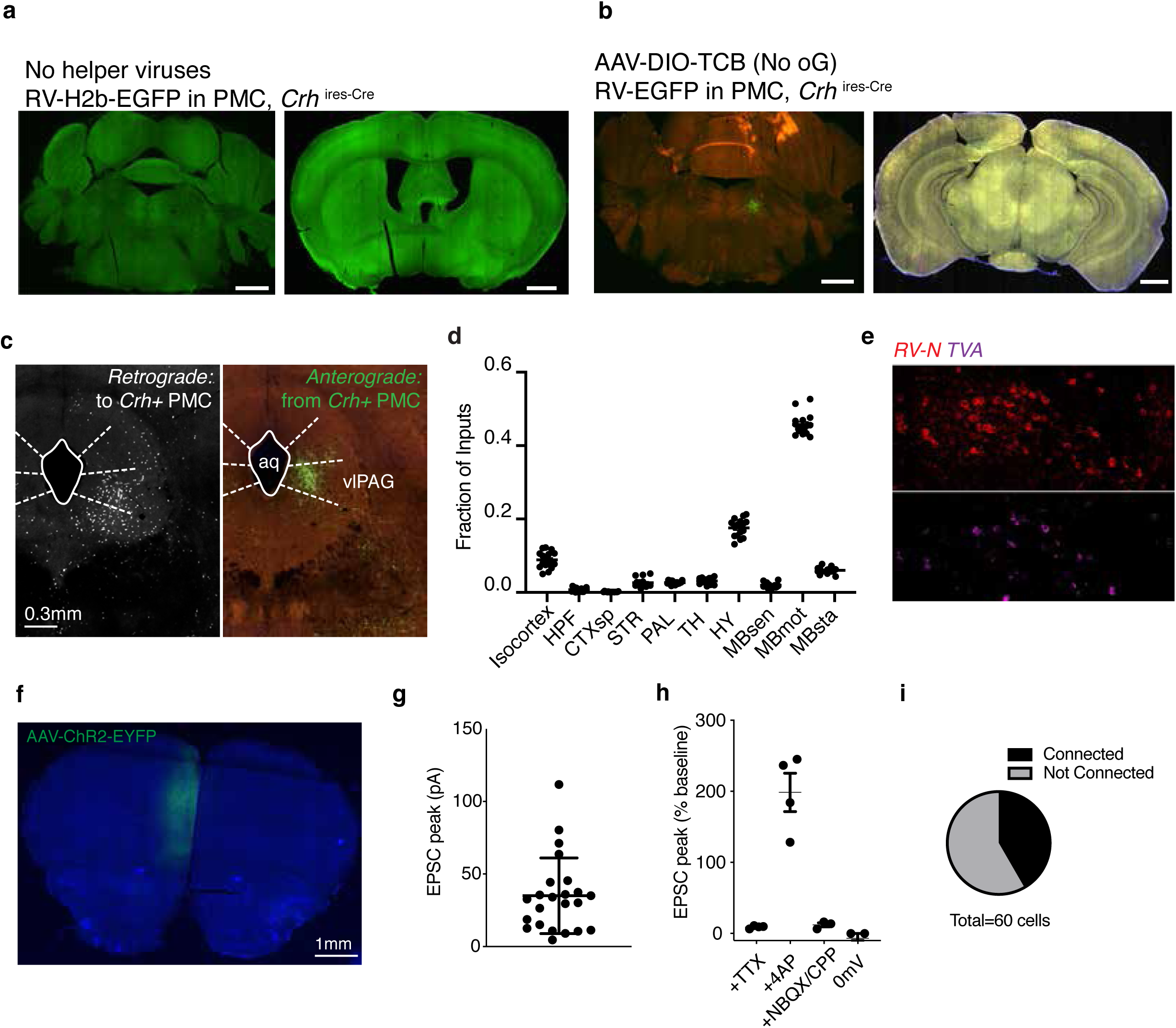
TCM in inbred laboratory strains. Related to figure 1. a) Comparison of micturition marks within a pair of male mice among 4 different laboratory strains. Mouse a always denotes partner with lower micturition marks (n=8 mice per strain) b) Comparison of micturition marks in paired and socially-isolated male mice among different strains after 1-hr TCM assay (n=8 pair-housed mice, n=7 socially-isolated mice).

**Supplementary Figure 2.**
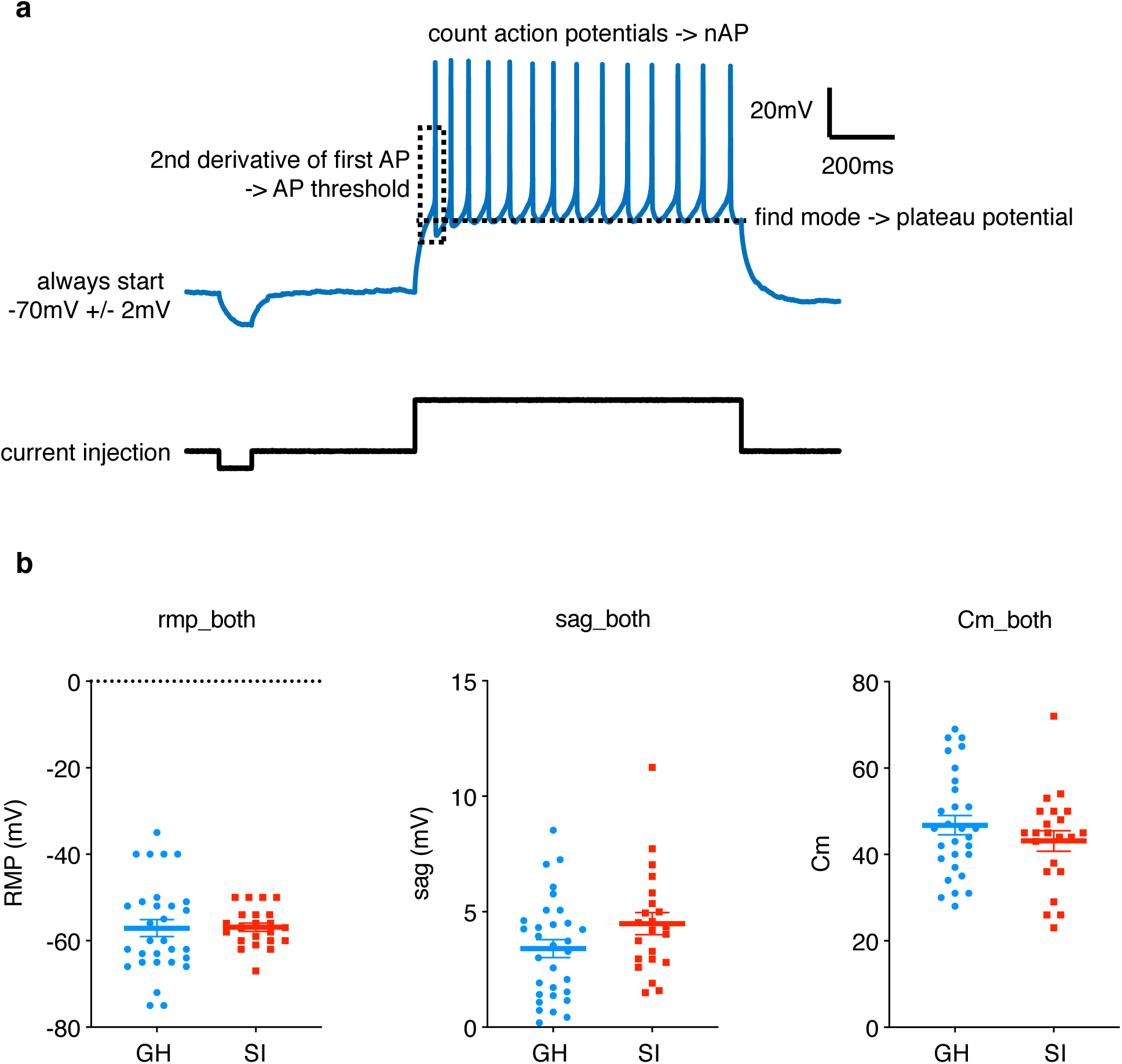
Supplementary materials for brain-wide rabies tracing. Related to figure 2. a) Helper virus injection was omitted, and RV-H2b-EGFP was injected into a *Crh*^ires-Cre^ mouse. No EGFP+ cells were observed in or around the PMC (left). No long-range EGFP+ input cells were present in the forebrain (right). b) AAV-DIO-RG was omitted from the helper AAV injection into a *Crh*^ires-Cre^ mouse, and RbV-EGFP was subsequently injected. Starter cell infection was observed in PMC (right), but no long-range EGFP+ cells were observed in the forebrain (left). Scale bars: 1 mm. c) PMC-PAG interconnectivity. Left: RV-h2b-EGFP labeled putative inputs to *Crh+* PMC in ventrolateral periaqueductal grey (vlPAG). Right: Anterograde tracing from *Crh+* PMC neurons (Allen Brain Institute) showing fibers from *Crh+* PMC neurons in vlPAG. d) Distributions of putative inputs to *Crh+* PMC neurons throughout the whole-brain. e) *In situ* Hybridization inside the PMC where *Rabies-N* mRNA (labels RV-infected cells) and *TVA* mRNA is visualized. *RV-N+* but *TVA-* cells are putative local presynaptic neurons to *Crh+* PMC starter neurons (*RV-N+/TVA+)* f) A coronal section of AAV-ChR2-EYFP infection in mPFC (DAPI in blue). g) Quantification of peak optogenetically evoked excitatory postsynaptic currents (EPSC) recorded from *Crh+* PMC neurons. h) Quantification of optogenetically-evoked EPSC currents normalized to baseline pre-drug EPSC amplitudes. i) Percentage of recorded *Crh+* PMC neurons that were responsive to optogenetic activation of mPFC inputs.

**Supplementary Figure 3.**
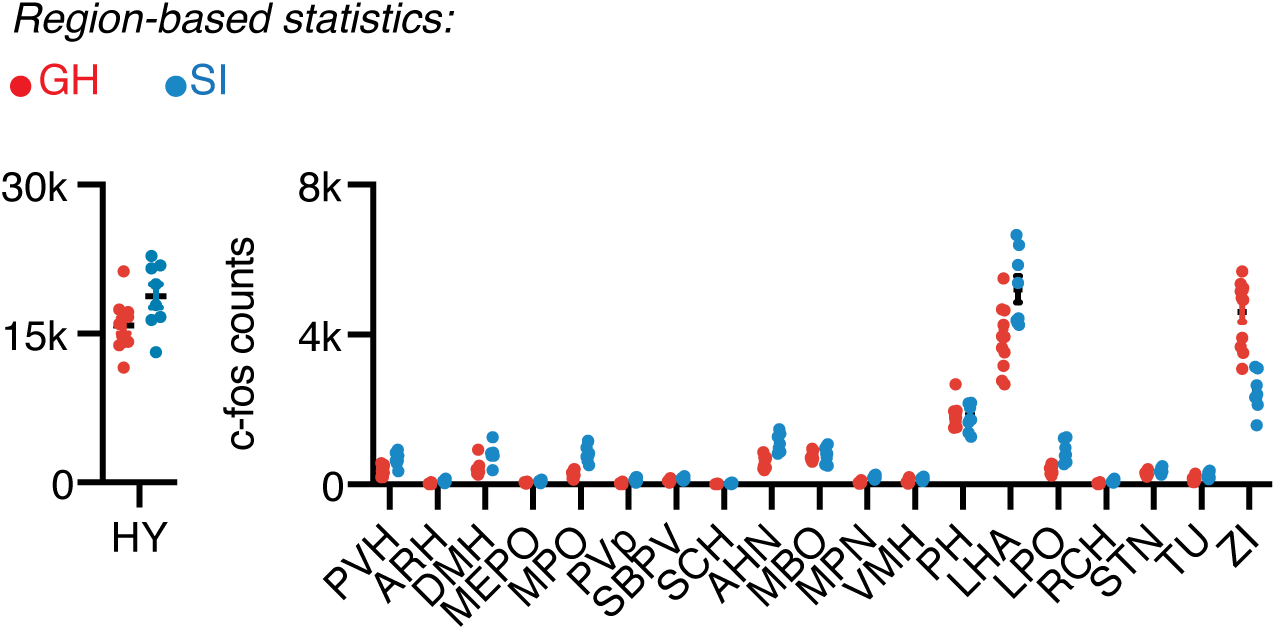
Analysis of current-clamp recordings of Crh+ PMC neurons and supplementary data. Related to figure 3. a) (Top) Example current-clamp recording trace with brief description of analysis methods, and (Bottom) corresponding current injection. b) (Left) Resting membrane potential (RMP) recordings in group-housed and socially-isolated males. (Middle) Sag potential recordings in group-housed and socially-isolated males. (Right) Membrane capacitance recordings in group-housed and socially-isolated males.

**Supplementary Figure 4.**
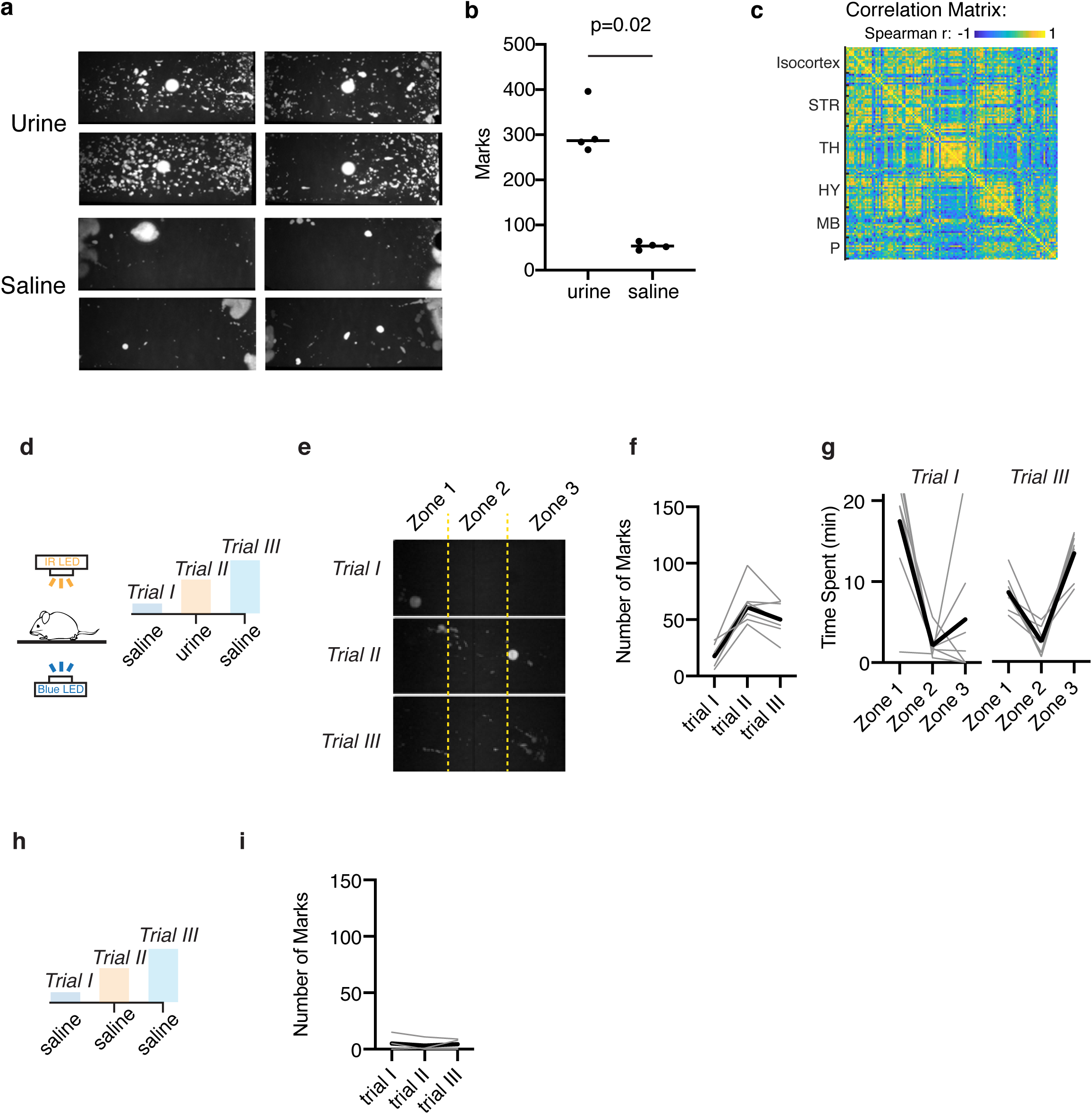
Supplementary materisl for brain-wide c-fos comparison between SI animals and GH animals. Related to figure 4. (left) c-fos immunolabeled cell count throughout hypothalamus (HY). (right)

**Supplementary Figure 5.**
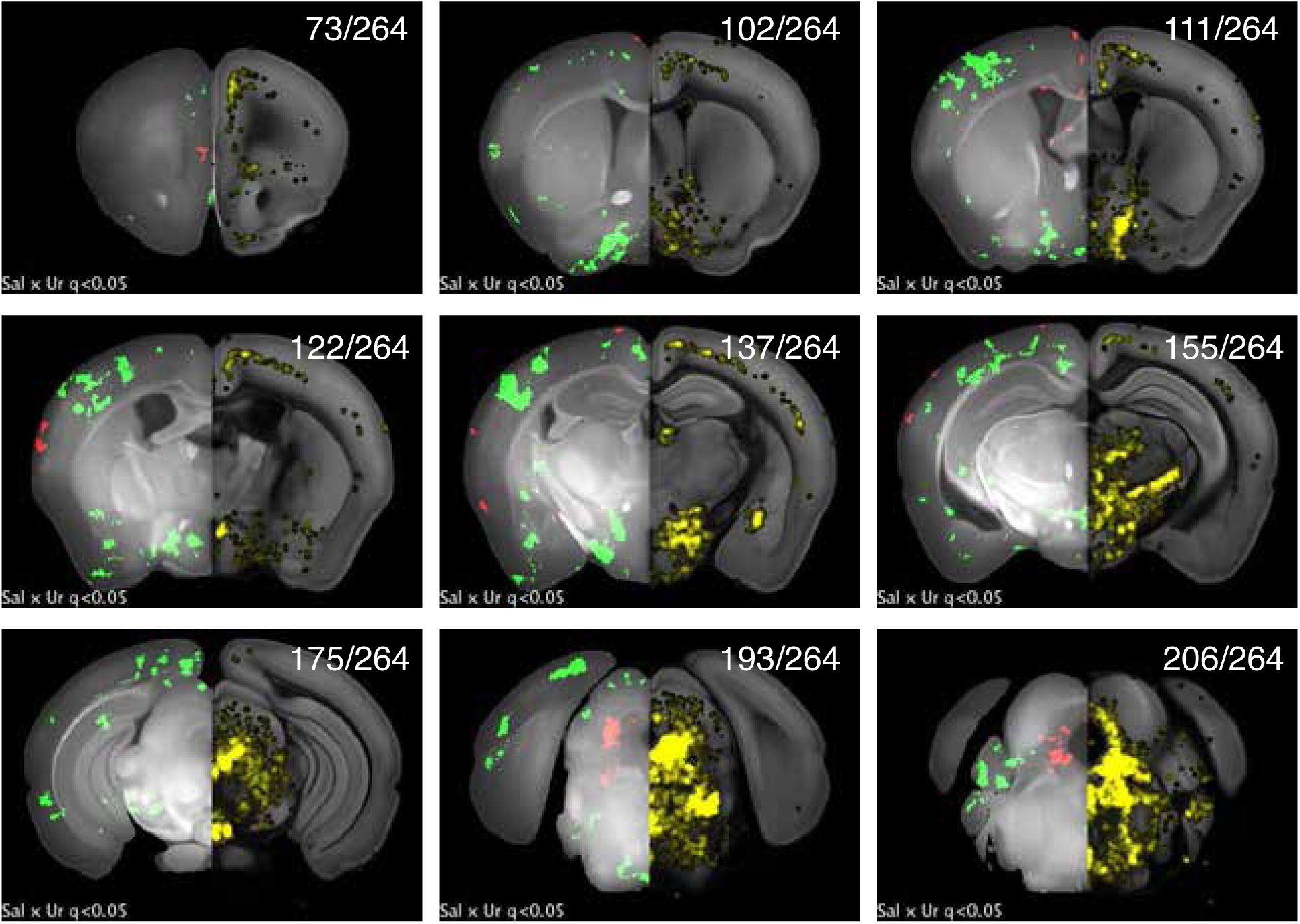
Supplementary materials for brain-wide c-fos comparison between SI animals exposed to urine and saline. Related to figure 5. a) TCM pattern from 4 Urine-exposed and 4 Saline-exposed mice used. b) Quantification of TCM between two groups (p=0.02, Mann-Whitney test). c) Brain-wide correlations for c-fos immunolabeling. d) Schematic of behavioral arena for real-time tracking of mouse position and micturition (left), and the trial design (right) where the saline (Trial I), male-urine (Trial II), and saline (Trial III) is applied in the arena. e) Example TCM patterns of a single mouse in Trials I, II, and III. Three zones for behavioral analysis are labeled with dotted yellow lines. f) Quantification of micturition marks in three trials (n=7 male mice, bold lines indicate the average). g) Quantification of the movement, where the time spent in each zone is calculated in Trials I (left) and III (right) (n=7 male mice, each trial = 20 minutes, bold lines indicate the average). h) Control experiment where a separate cohort of mice (n=4) was exposed to saline three trials in a row. i) Quantification of micturition marks in the control experiments.

**Supplementary Figure 6.**
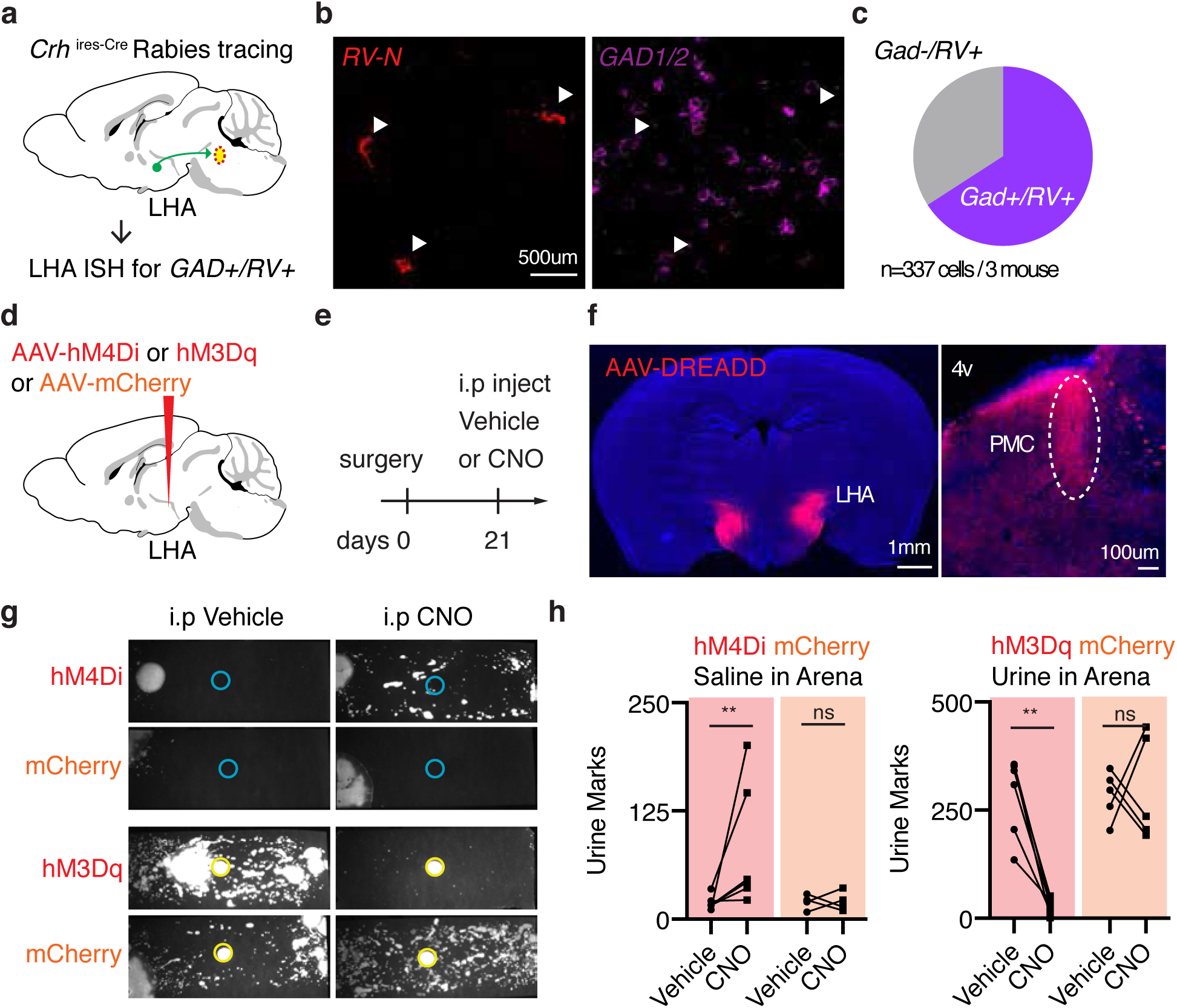
Related to figure 5. Statistical significance map of c-fos iDisco+ data (left) and average voxelated maps of rabies inputs of STPT (right) on the same coronal atlas based on ABA. Numbers at the top denotes its coronal location within 264 coronal sections spanning the whole-brain.

**Supplementary Table 1.** Lookup table for the abbreviated brain regions

| Acronym | Name |
| --- | --- |
| FRP | Frontal pole, cerebral cortex |
| MO | Somatomotor areas |
| SS | Somatosensory areas |
| ILA | Infralimbic area |
| GU | Gustatory areas |
| VISC | Visceral area |
| AUD | Auditory areas |
| VIS | Visual areas |
| ACA | Anterior cingulate area |
| PL | Prelimbic area |
| ORB | Orbital area |
| AI | Agranular insular area |
| RSP | Retrosplenial area |
| TEa | Temporal association areas |
| PERI | Perirhinal area |
| ECT | Ectorhinal area |
| CA | Ammon's horn |
| DG | Dentate gyrus |
| ENT | Entorhinal area |
| PAR | Parasubiculum |
| POST | Postsubiculum |
| PRE | Presubiculum |
| SUB | Subiculum |
| CLA | Clastrum |
| EP | Endopiriform nucleus |
| LA | Lateral amygdalar nucleus |
| BLA | Basolateral amygdalar nucleus |
| BMA | Basomedial amygdalar nucleus |
| PA | Posterior amygdalar nucleus |
| CP | Caudoputamen |
| ACB | Nucleus accumbens |
| FS | Fundus of striatum |
| OT | Olfactory tubercle |
| LS | Lateral septal nucleus |
| SF | Septofimbrial nucleus |
| AAA | Anterior amygdalar area |
| CEA | Central amygdalar nucleus |
| IA | Intercalated amygdalar nucleus |
| MEA | Medial amygdalar nucleus |
| GPe | Globus pallidus, external segment |
| GPI | Globus pallidus, internal segment |
| SI | Substantia innominata |
| MA | Magnocellular nucleus |
| MSC | Medial septal complex |
| TRS | Triangular nucleus of septum |
| BST | Bed nuclei of the stria terminalis |
| VAL | Ventral anterior-lateral complex of the thalamus |
| VM | Ventral medial nucleus of the thalamus |
| VP | Ventral posterior complex of the thalamus |
| SPF | Subparafascicular nucleus |
| MG | Medial geniculate complex |
| LGd | Dorsal part of the lateral geniculate complex |
| LP | Lateral posterior nucleus of the thalamus |
| PO | Posterior complex of the thalamus |
| POL | Posterior limiting nucleus of the thalamus |
| AV | Anteroventral nucleus of thalamus |
| AM | Anteromedial nucleus |
| AD | Anterodorsal nucleus |
| LD | Lateral dorsal nucleus of thalamus |
| IMD | Intermediodorsal nucleus of the thalamus |
| MD | Mediodorsal nucleus of thalamus |
| SMT | Submedial nucleus of the thalamus |
| PVT | Paraventricular nucleus of the thalamus |
| PT | Parataenial nucleus |
| RE | Nucleus of reunions |
| CM | Central medial nucleus of the thalamus |
| PCN | Paracentral nucleus |
| CL | Central lateral nucleus of the thalamus |
| PF | Parafascicular nucleus |
| RT | Reticular nucleus of the thalamus |
| LGv | Ventral part of the lateral geniculate complex |
| MH | Medial habenula |
| LH | Lateral habenula |
| PVH | Paraventricular hypothalamic nucleus |
| ARH | Arcuate hypothalamic nucleus |
| DMH | Dorsomedial nucleus of the hypothalamus |
| MEPO | Median preoptic nucleus |
| MPO | Medial preoptic area |
| PVp | Periventricular hypothalamic nucleus, posterior part |
| SBPV | Subparaventricular zone |
| SCH | Suprachiasmatic nucleus |
| AHN | Anterior hypothalamic nucleus |
| MBO | Mammillary body |
| MPN | Medial preoptic nucleus |
| VMH | Ventromedial hypothalamic nucleus |
| PH | Posterior hypothalamic nucleus |
| LHA | Lateral hypothalamic area |
| LPO | Lateral preoptic area |
| RCH | Retrochiasmatic area |
| STN | Subthalamic nucleus |
| TU | Tuberal nucleus |
| ZI | Zona incerta |
| SCs | Superior colliculus, sensory related |
| IC | Inferior colliculus |
| SNr | Substantia nigra, reticular part |
| VTA | Ventral tegmental area |
| RR | Midbrain reticular nucleus, retrorubral area |
| MRN | Midbrain reticular nucleus |
| SCm | Superior colliculus, motor related |
| PAG | Periaqueductal gray |
| CUN | Cuneiform nucleus |
| RN | Red nucleus |
| SNc | Substantia nigra, compact part |
| PPN | Pedunculo pontine nucleus |
| RAmb | Midbrain raphé nuclei |
| NLL | Nucleus of the lateral lemniscus |
| PSV | Principal sensory nucleus of the trigeminal |
| PB | Parabrachial nucleus |
| SOC | Superior olivary complex |
| B | Barrington's nucleus |
| PCG | Pontine central gray |
| PG | Pontine gray |
| PRNc | Pontine reticular nucleus, caudal part |
| SUT | Supratrigeminal nucleus |
| TRN | Tegmental reticular nucleus |
| V | Motor nucleus of trigeminal |
| CS | Superior central nucleus raphé |
| LC | Locus ceruleus |
| LDT | Laterodorsal tegmental nucleus |
| NI | Nucleus incertus |
| PRNr | Pontine reticular nucleus |

**Supplementary Table 2.** Summary region-based statistics table from brain-wide distribution of putative inputs to *Crh+* PMC neurons, normalized by total number of forebrain RV+ input cells (n=6 mice for GH-male, n=4 mice for SI-male, n=6 mice for GH-Female. FDR=0.05 for q-value)

| ROIs | GH-male mean | GH-female mean | SI-male mean | p (Male vs Female) | p (GH Male vs SI Male) | p (GH-Female vs SI-Male) | q (GH-Male vs GH-Female) | q (GH-Male vs SI-Male) | q (GH-Female vs SI-Male) |
| --- | --- | --- | --- | --- | --- | --- | --- | --- | --- |
| FRP | 0.0005 | 0.0005 | 0.0002 | 0.8182 | 0.0095 | 0.3923 | 1.0000 | 0.6968 | 0.9505 |
| MO | 0.0314 | 0.0382 | 0.0346 | 0.5887 | 0.9143 | 0.7619 | 1.0000 | 1.0000 | 1.0000 |
| SS | 0.0203 | 0.0202 | 0.0136 | 0.9372 | 0.2571 | 0.1714 | 1.0000 | 0.8410 | 0.8803 |
| ILA | 0.0035 | 0.0035 | 0.0019 | 0.9372 | 0.1714 | 0.1714 | 1.0000 | 0.7643 | 0.8803 |
| GU | 0.0012 | 0.0013 | 0.0015 | 0.6991 | 1.0000 | 0.7619 | 1.0000 | 1.0000 | 1.0000 |
| VISC | 0.0006 | 0.0005 | 0.0011 | 0.4848 | 0.7619 | 0.9143 | 1.0000 | 0.9962 | 1.0000 |
| AUD | 0.0035 | 0.0033 | 0.0024 | 0.8182 | 0.3524 | 0.3524 | 1.0000 | 0.8602 | 0.8803 |
| VIS | 0.0048 | 0.0051 | 0.0034 | 0.8182 | 0.3524 | 0.2571 | 1.0000 | 0.8602 | 0.8803 |
| ACA | 0.0109 | 0.0110 | 0.0068 | 0.9372 | 0.3524 | 0.1714 | 1.0000 | 0.8602 | 0.8803 |
| PL | 0.0028 | 0.0025 | 0.0018 | 0.8182 | 0.0667 | 0.3524 | 1.0000 | 0.6968 | 0.8803 |
| ORB | 0.0082 | 0.0085 | 0.0068 | 0.8182 | 0.7619 | 0.3524 | 1.0000 | 0.9962 | 0.8803 |
| AI | 0.0024 | 0.0025 | 0.0029 | 0.9372 | 0.7619 | 0.7619 | 1.0000 | 0.9962 | 1.0000 |
| RSP | 0.0085 | 0.0096 | 0.0085 | 0.8182 | 0.3524 | 0.2571 | 1.0000 | 0.8602 | 0.8803 |
| TEa | 0.0012 | 0.0012 | 0.0013 | 0.6991 | 0.6095 | 0.9143 | 1.0000 | 0.9521 | 1.0000 |
| PERI | 0.0006 | 0.0005 | 0.0008 | 0.3939 | 0.6095 | 0.4762 | 1.0000 | 0.9521 | 1.0000 |
| ECT | 0.0010 | 0.0011 | 0.0016 | 0.8182 | 0.7619 | 1.0000 | 1.0000 | 0.9962 | 1.0000 |
| CA | 0.0015 | 0.0008 | 0.0004 | 0.3095 | 0.0667 | 0.3524 | 1.0000 | 0.6968 | 0.8803 |
| DG | 0.0025 | 0.0012 | 0.0006 | 0.1320 | 0.0190 | 0.1714 | 1.0000 | 0.6968 | 0.8803 |
| ENT | 0.0026 | 0.0022 | 0.0021 | 0.9372 | 0.3524 | 0.7619 | 1.0000 | 0.8602 | 1.0000 |
| PAR | 0.0003 | 0.0003 | 0.0000 | 0.6991 | 0.0114 | 0.0307 | 1.0000 | 0.6968 | 0.8803 |
| POST | 0.0003 | 0.0001 | 0.0001 | 0.0931 | 0.0666 | 0.2714 | 1.0000 | 0.6968 | 0.8803 |
| PRE | 0.0002 | 0.0002 | 0.0000 | 0.6868 | 0.0477 | 0.1393 | 1.0000 | 0.6968 | 0.8803 |
| SUB | 0.0010 | 0.0006 | 0.0002 | 0.4848 | 0.0381 | 0.0190 | 1.0000 | 0.6968 | 0.8803 |
| CLA | 0.0007 | 0.0009 | 0.0007 | 0.5887 | 1.0000 | 0.6095 | 1.0000 | 1.0000 | 1.0000 |
| EP | 0.0001 | 0.0003 | 0.0005 | 0.0649 | 0.2571 | 0.7619 | 1.0000 | 0.8410 | 1.0000 |
| LA | 0.0001 | 0.0001 | 0.0000 | 0.8089 | 0.1236 | 0.2110 | 1.0000 | 0.7487 | 0.8803 |
| BLA | 0.0001 | 0.0002 | 0.0001 | 0.9372 | 0.2352 | 0.2714 | 1.0000 | 0.8410 | 0.8803 |
| BMA | 0.0001 | 0.0002 | 0.0002 | 0.2539 | 1.0000 | 0.5895 | 1.0000 | 1.0000 | 1.0000 |
| PA | 0.0000 | 0.0000 | 0.0000 | 0.4460 | 0.1491 | 0.5403 | 1.0000 | 0.7643 | 1.0000 |
| CP | 0.0038 | 0.0050 | 0.0023 | 0.5887 | 0.2571 | 0.1714 | 1.0000 | 0.8410 | 0.8803 |
| ACB | 0.0015 | 0.0015 | 0.0012 | 0.9372 | 0.7619 | 0.2571 | 1.0000 | 0.9962 | 0.8803 |
| FS | 0.0003 | 0.0003 | 0.0006 | 0.9372 | 0.0381 | 0.0381 | 1.0000 | 0.6968 | 0.8803 |
| OT | 0.0014 | 0.0012 | 0.0005 | 0.5887 | 0.0190 | 0.3524 | 1.0000 | 0.6968 | 0.8803 |
| LS | 0.0011 | 0.0019 | 0.0003 | 0.8182 | 0.0422 | 0.0422 | 1.0000 | 0.6968 | 0.8803 |
| SF | 0.0001 | 0.0001 | 0.0000 | 0.6868 | 0.5824 | 0.3314 | 1.0000 | 0.9521 | 0.8803 |
| AAA | 0.0008 | 0.0008 | 0.0008 | 0.8182 | 0.7619 | 0.9143 | 1.0000 | 0.9962 | 1.0000 |
| CEA | 0.0145 | 0.0163 | 0.0190 | 0.5887 | 0.4762 | 0.7619 | 1.0000 | 0.9349 | 1.0000 |
| IA | 0.0000 | 0.0000 | 0.0000 | 0.4942 | 0.4712 | 0.8102 | 1.0000 | 0.9349 | 1.0000 |
| MEA | 0.0008 | 0.0007 | 0.0011 | 0.6991 | 1.0000 | 1.0000 | 1.0000 | 1.0000 | 1.0000 |
| GPe | 0.0005 | 0.0005 | 0.0005 | 0.6991 | 1.0000 | 0.9143 | 1.0000 | 1.0000 | 1.0000 |
| GPi | 0.0001 | 0.0002 | 0.0002 | 0.0904 | 0.2352 | 1.0000 | 1.0000 | 0.8410 | 1.0000 |
| SI | 0.0051 | 0.0051 | 0.0059 | 0.9372 | 0.1143 | 0.1714 | 1.0000 | 0.7411 | 0.8803 |
| MA | 0.0008 | 0.0010 | 0.0011 | 0.2403 | 0.1143 | 0.2571 | 1.0000 | 0.7411 | 0.8803 |
| MSC | 0.0020 | 0.0028 | 0.0018 | 0.3939 | 0.7619 | 0.2571 | 1.0000 | 0.9962 | 0.8803 |
| TRS | 0.0001 | 0.0001 | 0.0000 | 0.8182 | 0.0477 | 0.0131 | 1.0000 | 0.6968 | 0.8803 |
| BST | 0.0197 | 0.0191 | 0.0186 | 0.9372 | 0.9143 | 0.9143 | 1.0000 | 1.0000 | 1.0000 |
| VAL | 0.0001 | 0.0001 | 0.0002 | 0.8705 | 0.4500 | 0.3524 | 1.0000 | 0.9349 | 0.8803 |
| VM | 0.0033 | 0.0034 | 0.0029 | 0.9372 | 0.7619 | 0.3524 | 1.0000 | 0.9962 | 0.8803 |
| VP | 0.0003 | 0.0007 | 0.0004 | 0.1275 | 0.4542 | 0.2571 | 1.0000 | 0.9349 | 0.8803 |
| SPF | 0.0011 | 0.0010 | 0.0013 | 0.8182 | 0.7619 | 0.6095 | 1.0000 | 0.9962 | 1.0000 |
| MG | 0.0001 | 0.0001 | 0.0001 | 1.0000 | 1.0000 | 0.9143 | 1.0000 | 1.0000 | 1.0000 |
| LGd | 0.0000 | 0.0000 | 0.0000 | 0.4047 | 0.5403 | 1.0000 | 1.0000 | 0.9521 | 1.0000 |
| LP | 0.0001 | 0.0001 | 0.0000 | 0.7483 | 0.0307 | 0.0307 | 1.0000 | 0.6968 | 0.8803 |
| PO | 0.0003 | 0.0004 | 0.0002 | 0.8182 | 0.1714 | 0.1714 | 1.0000 | 0.7643 | 0.8803 |
| POL | 0.0002 | 0.0002 | 0.0002 | 0.9372 | 0.7619 | 0.6095 | 1.0000 | 0.9962 | 1.0000 |
| AV | 0.0000 | 0.0000 | 0.0000 | 1.0000 | 1.0000 | 1.0000 | 1.0000 | 1.0000 | 1.0000 |
| AM | 0.0001 | 0.0000 | 0.0000 | 0.4620 | 0.6940 | 1.0000 | 1.0000 | 0.9962 | 1.0000 |
| AD | 0.0000 | 0.0000 | 0.0000 | 0.4047 | 0.5120 | 0.0942 | 1.0000 | 0.9521 | 0.8803 |
| LD | 0.0000 | 0.0000 | 0.0001 | 0.9289 | 0.5482 | 0.5696 | 1.0000 | 0.9521 | 1.0000 |
| IMD | 0.0003 | 0.0002 | 0.0002 | 0.8182 | 0.6689 | 0.6095 | 1.0000 | 0.9962 | 1.0000 |
| MD | 0.0002 | 0.0001 | 0.0004 | 0.1994 | 0.1143 | 0.1143 | 1.0000 | 0.7411 | 0.8803 |
| SMT | 0.0000 | 0.0000 | 0.0000 | 0.4047 | 1.0000 | 0.5403 | 1.0000 | 1.0000 | 1.0000 |
| PVT | 0.0009 | 0.0009 | 0.0008 | 0.9372 | 1.0000 | 1.0000 | 1.0000 | 1.0000 | 1.0000 |
| PT | 0.0000 | 0.0000 | 0.0000 | 1.0000 | 0.7609 | 0.7609 | 1.0000 | 0.9962 | 1.0000 |
| RE | 0.0003 | 0.0002 | 0.0003 | 0.3358 | 1.0000 | 0.9143 | 1.0000 | 1.0000 | 1.0000 |
| CM | 0.0000 | 0.0001 | 0.0001 | 0.7750 | 0.5482 | 1.0000 | 1.0000 | 0.9521 | 1.0000 |
| PCN | 0.0000 | 0.0000 | 0.0000 | 0.4047 | 1.0000 | 0.5403 | 1.0000 | 1.0000 | 1.0000 |
| CL | 0.0000 | 0.0000 | 0.0000 | 0.7526 | 0.7186 | 0.5120 | 1.0000 | 0.9962 | 1.0000 |
| PF | 0.0015 | 0.0015 | 0.0015 | 0.9372 | 0.9143 | 1.0000 | 1.0000 | 1.0000 | 1.0000 |
| RT | 0.0004 | 0.0003 | 0.0002 | 0.5887 | 0.9143 | 0.2571 | 1.0000 | 1.0000 | 0.8803 |
| LGv | 0.0006 | 0.0004 | 0.0005 | 0.5887 | 0.7619 | 0.9143 | 1.0000 | 0.9962 | 1.0000 |
| MH | 0.0021 | 0.0021 | 0.0013 | 1.0000 | 0.2571 | 0.3524 | 1.0000 | 0.8410 | 0.8803 |
| LH | 0.0082 | 0.0079 | 0.0081 | 0.8182 | 1.0000 | 1.0000 | 1.0000 | 1.0000 | 1.0000 |
| PVH | 0.0035 | 0.0024 | 0.0032 | 0.3939 | 0.9143 | 0.6095 | 1.0000 | 1.0000 | 1.0000 |
| ARH | 0.0025 | 0.0017 | 0.0013 | 0.1797 | 0.0667 | 0.6095 | 1.0000 | 0.6968 | 1.0000 |
| DMH | 0.0122 | 0.0104 | 0.0088 | 0.3095 | 0.1143 | 0.4762 | 1.0000 | 0.7411 | 1.0000 |
| MEP | 0.0013 | 0.0012 | 0.0008 | 0.8182 | 0.3524 | 0.1714 | 1.0000 | 0.8602 | 0.8803 |
| O |  |  |  |  |  |  |  |  |  |
| MPO | 0.0091 | 0.0093 | 0.0069 | 0.9372 | 0.2571 | 0.2571 | 1.0000 | 0.8410 | 0.8803 |
| PVp | 0.0009 | 0.0012 | 0.0005 | 0.4848 | 0.0667 | 0.0381 | 1.0000 | 0.6968 | 0.8803 |
| SBPV | 0.0013 | 0.0014 | 0.0013 | 0.8182 | 1.0000 | 0.9143 | 1.0000 | 1.0000 | 1.0000 |
| SCH | 0.0000 | 0.0002 | 0.0000 | 0.1581 | 0.6940 | 0.1236 | 1.0000 | 0.9962 | 0.8803 |
| AHN | 0.0021 | 0.0020 | 0.0021 | 0.6991 | 1.0000 | 0.9143 | 1.0000 | 1.0000 | 1.0000 |
| MBO | 0.0056 | 0.0056 | 0.0044 | 0.8182 | 0.1714 | 0.4762 | 1.0000 | 0.7643 | 1.0000 |
| MPN | 0.0016 | 0.0023 | 0.0021 | 0.3095 | 0.6095 | 0.9143 | 1.0000 | 0.9521 | 1.0000 |
| VMH | 0.0025 | 0.0014 | 0.0020 | 0.0931 | 0.9143 | 0.2571 | 1.0000 | 1.0000 | 0.8803 |
| PH | 0.0244 | 0.0207 | 0.0211 | 0.3939 | 0.6095 | 0.9143 | 1.0000 | 0.9521 | 1.0000 |
| LHA | 0.0608 | 0.0544 | 0.0533 | 0.3939 | 0.2571 | 0.7619 | 1.0000 | 0.8410 | 1.0000 |
| LPO | 0.0066 | 0.0060 | 0.0069 | 0.8182 | 0.9143 | 0.6095 | 1.0000 | 1.0000 | 1.0000 |
| RCH | 0.0005 | 0.0005 | 0.0002 | 1.0000 | 0.2850 | 0.2571 | 1.0000 | 0.8410 | 0.8803 |
| STN | 0.0018 | 0.0016 | 0.0023 | 0.8182 | 0.2571 | 0.1714 | 1.0000 | 0.8410 | 0.8803 |
| TU | 0.0030 | 0.0020 | 0.0017 | 0.3939 | 0.2571 | 0.7619 | 1.0000 | 0.8410 | 1.0000 |
| ZI | 0.0296 | 0.0320 | 0.0343 | 0.8182 | 0.4762 | 1.0000 | 1.0000 | 0.9349 | 1.0000 |
| SCs | 0.0009 | 0.0007 | 0.0009 | 0.8182 | 1.0000 | 0.9143 | 1.0000 | 1.0000 | 1.0000 |
| IC | 0.0075 | 0.0094 | 0.0124 | 0.6991 | 1.0000 | 0.9143 | 1.0000 | 1.0000 | 1.0000 |
| SNr | 0.0076 | 0.0080 | 0.0132 | 0.8182 | 0.1143 | 0.0667 | 1.0000 | 0.7411 | 0.8803 |
| VTA | 0.0092 | 0.0091 | 0.0106 | 1.0000 | 0.2571 | 0.1143 | 1.0000 | 0.8410 | 0.8803 |
| RR | 0.0123 | 0.0116 | 0.0177 | 0.9372 | 0.1143 | 0.0381 | 1.0000 | 0.7411 | 0.8803 |
| MRN | 0.0823 | 0.0936 | 0.1053 | 0.1797 | 0.0095 | 0.1714 | 1.0000 | 0.6968 | 0.8803 |
| SCm | 0.0519 | 0.0645 | 0.0650 | 0.3939 | 0.0381 | 0.7619 | 1.0000 | 0.6968 | 1.0000 |
| PAG | 0.2544 | 0.2350 | 0.2466 | 0.3095 | 0.9143 | 0.4762 | 1.0000 | 1.0000 | 1.0000 |
| CUN | 0.0041 | 0.0040 | 0.0040 | 0.8182 | 0.7619 | 1.0000 | 1.0000 | 0.9962 | 1.0000 |
| RN | 0.0013 | 0.0028 | 0.0015 | 0.8182 | 0.7619 | 0.6095 | 1.0000 | 0.9962 | 1.0000 |
| SNc | 0.0044 | 0.0045 | 0.0074 | 0.9372 | 0.1714 | 0.1143 | 1.0000 | 0.7643 | 0.8803 |
| PPN | 0.0167 | 0.0168 | 0.0184 | 0.5887 | 0.6095 | 0.3524 | 1.0000 | 0.9521 | 0.8803 |
| RAm<br>b | 0.0334 | 0.0360 | 0.0310 | 0.5887 | 0.7619 | 0.7619 | 1.0000 | 0.9962 | 1.0000 |

**Supplementary Table 3.** Summary region-based statistics table from brain-wide c-fos count comparison between single-housed males and pair-housed males (n=8 for single-housed and n=11 for group-housed biologically independent samples. FDR=0.05 for q-value)

| ROIs | mean.single | std.single | mean.pair | std.pair | p-value | q-value |
| --- | --- | --- | --- | --- | --- | --- |
| FRP | 8 | 7.78276484 | 3.63636364 | 2.69342634 | 0.05167725 | 0.09867253 |
| MO | 9474.5 | 2310.68734 | 2138.54545 | 606.273925 | 6.07E-34 | 2.80E-32 |
| SS | 37380.125 | 9731.39041 | 9930.27273 | 2511.03346 | 2.79E-31 | 1.08E-29 |
| ILA | 1508.125 | 374.516617 | 229.181818 | 228.420147 | 9.95E-10 | 3.99E-09 |
| GU | 796.25 | 273.602814 | 193.363636 | 65.6205345 | 1.91E-20 | 2.31E-19 |
| VISC | 1879.625 | 445.7568 | 449.454545 | 210.895881 | 3.06E-17 | 2.62E-16 |
| AUD | 25722.75 | 3659.13837 | 11576.3636 | 2015.48149 | 6.98E-30 | 2.09E-28 |
| VIS | 51038 | 6992.74097 | 19436.2727 | 6396.2222 | 1.11E-16 | 8.93E-16 |
| ACA | 3910.25 | 769.536363 | 1090.45455 | 346.650072 | 3.14E-28 | 8.35E-27 |
| PL | 840 | 134.437452 | 63.2727273 | 50.3251248 | 6.49E-19 | 6.59E-18 |
| ORB | 1019.875 | 335.884095 | 259.909091 | 122.861267 | 6.00E-13 | 3.16E-12 |
| AI | 3308.5 | 1070.91896 | 900.363636 | 283.193317 | 1.41E-18 | 1.41E-17 |
| RSP | 14287.25 | 2018.48527 | 5593.18182 | 1211.34304 | 2.37E-29 | 6.61E-28 |
| TEa | 13003.125 | 2030.78313 | 6484.45455 | 1275.57739 | 5.84E-19 | 6.03E-18 |
| PERI | 4940.125 | 671.090677 | 3336.09091 | 625.697284 | 3.90E-07 | 1.23E-06 |
| ECT | 13587.375 | 2101.77938 | 7439.81818 | 1423.85363 | 8.73E-15 | 5.63E-14 |
| CA | 12968.75 | 1300.50372 | 9122.72727 | 741.27459 | 6.25E-19 | 6.40E-18 |
| DG | 5504.875 | 565.877684 | 5507.27273 | 926.693271 | 0.99470159 | 1 |
| ENT | 35314.375 | 4604.23212 | 22176.8182 | 1896.29775 | 5.52E-23 | 8.39E-22 |
| PAR | 3847.875 | 732.782259 | 2509.72727 | 451.256488 | 1.64E-07 | 5.35E-07 |
| POST | 2674.75 | 305.771039 | 1732.63636 | 403.768566 | 3.19E-07 | 1.01E-06 |
| PRE | 4836.375 | 513.651894 | 3205 | 333.433652 | 9.41E-19 | 9.48E-18 |
| SUB | 13467.625 | 1167.36112 | 8723.18182 | 802.483373 | 9.68E-27 | 2.11E-25 |
| CLA | 1719.5 | 190.115003 | 631.727273 | 184.832406 | 3.36E-21 | 4.49E-20 |
| EP | 4412.125 | 766.91412 | 2492.90909 | 390.688995 | 1.72E-14 | 1.06E-13 |
| LA | 823 | 409.0358 | 529.636364 | 206.877874 | 0.01471798 | 0.02990993 |
| BLA | 2936.625 | 524.318861 | 1874.36364 | 624.916838 | 0.00011272 | 0.00027983 |
| BMA | 2534.625 | 501.47324 | 1289.27273 | 247.252539 | 1.76E-14 | 1.07E-13 |
| PA | 683.625 | 96.6376701 | 429.545455 | 112.034248 | 2.01E-06 | 5.89E-06 |
| CP | 9276.875 | 1485.62824 | 6820.27273 | 1293.6304 | 9.33E-05 | 0.00023359 |
| ACB | 1781.5 | 401.607129 | 705.818182 | 232.304033 | 3.55E-13 | 1.92E-12 |
| FS | 231.625 | 52.5627176 | 129 | 77.9102047 | 0.00113381 | 0.00261933 |
| OT | 979.125 | 231.175529 | 201.909091 | 95.0373132 | 3.44E-19 | 3.72E-18 |
| LS | 2439.625 | 987.57422 | 816.727273 | 268.756057 | 6.60E-11 | 2.93E-10 |
| SF | 582.25 | 364.854864 | 409 | 221.142036 | 0.38191238 | 0.627278 |
| AAA | 749 | 241.240248 | 368.636364 | 98.5629471 | 9.00E-08 | 3.01E-07 |
| CEA | 1434.5 | 276.400331 | 853.363636 | 454.129117 | 0.00078435 | 0.00183678 |
| IA | 472.125 | 88.2778689 | 219.272727 | 63.909453 | 4.62E-12 | 2.24E-11 |
| MEA | 2769.375 | 526.617763 | 1497.18182 | 399.15406 | 7.74E-09 | 2.85E-08 |
| GPe | 320.625 | 61.2394306 | 506 | 139.293934 | 2.31E-05 | 6.08E-05 |
| GPI | 97.5 | 23.4520788 | 262.363636 | 101.607355 | 3.94E-08 | 1.36E-07 |
| SI | 972.25 | 236.076713 | 607.363636 | 120.939053 | 1.35E-06 | 4.05E-06 |
| MA | 151.75 | 99.7836946 | 83 | 36.1718122 | 0.01405278 | 0.02860658 |
| MSC | 706.75 | 203.457788 | 330.909091 | 73.9911543 | 4.32E-12 | 2.11E-11 |
| TRS | 208.5 | 157.024111 | 165.545455 | 96.9859409 | 0.56032455 | 0.88166553 |
| BST | 1777.5 | 588.570908 | 643.272727 | 161.744299 | 1.45E-14 | 9.08E-14 |
| VAL | 153 | 142.652425 | 435.181818 | 209.302087 | 0.00121085 | 0.00278125 |
| VM | 272.5 | 94.1123948 | 406.454545 | 104.327718 | 0.00284802 | 0.00622967 |
| VP | 180.875 | 143.047882 | 2103 | 755.4727 | 1.49E-23 | 2.56E-22 |
| SPF | 516.625 | 102.611525 | 678.363636 | 92.1089276 | 8.99E-05 | 0.00022637 |
| MG | 787.125 | 266.58284 | 684.909091 | 167.700599 | 0.24454795 | 0.41947496 |
| LGd | 79.125 | 78.4409469 | 362.727273 | 133.032395 | 7.37E-10 | 2.97E-09 |
| LP | 338.875 | 306.076292 | 326.181818 | 77.4181092 | 0.86473098 | 1 |
| PO | 155.75 | 188.54916 | 404.818182 | 123.316518 | 0.00056425 | 0.0013344 |
| POL | 120.375 | 33.1961164 | 227.181818 | 39.7764206 | 1.11E-11 | 5.13E-11 |
| AV | 522.125 | 470.06639 | 277.272727 | 182.168653 | 0.07972087 | 0.14773619 |
| AM | 674.5 | 413.222873 | 278.636364 | 104.290242 | 0.00018877 | 0.00046381 |
| AD | 216.5 | 153.216933 | 123.818182 | 57.3982895 | 0.07140528 | 0.13273632 |
| LD | 148.625 | 83.4333566 | 297.818182 | 68.5285607 | 0.00023616 | 0.00057552 |
| IMD | 306 | 125.434103 | 260.181818 | 132.393971 | 0.45142635 | 0.7334149 |
| MD | 773 | 332.059805 | 663.545455 | 250.826778 | 0.4549281 | 0.73713482 |
| SMT | 31 | 25.3940375 | 29.8181818 | 10.1766221 | 0.87901879 | 1 |
| PVT | 2249.75 | 647.642924 | 1957.18182 | 667.246404 | 0.40805965 | 0.66748093 |
| PT | 521.625 | 205.16261 | 372.727273 | 121.59037 | 0.05233662 | 0.09960574 |
| RE | 1361.5 | 517.744283 | 1195.09091 | 220.106999 | 0.33289763 | 0.55359815 |
| CM | 447.375 | 115.881636 | 328.090909 | 55.9490028 | 0.00109979 | 0.00255057 |
| PCN | 55.625 | 35.7648571 | 37.4545455 | 10.6991928 | 0.06795918 | 0.12688431 |
| CL | 149.25 | 82.5915077 | 86.4545455 | 52.236699 | 0.04255162 | 0.0828237 |
| PF | 318.375 | 179.288226 | 270 | 167.512388 | 0.53914667 | 0.85281907 |
| RT | 466 | 175.730313 | 3061.63636 | 1065.49935 | 1.37E-27 | 3.35E-26 |
| LGv | 678.875 | 112.583223 | 864.090909 | 220.432055 | 0.01785526 | 0.03586006 |
| MH | 122 | 45.9409559 | 200.818182 | 121.505406 | 0.0317276 | 0.06267117 |
| LH | 267.75 | 103.673595 | 293 | 116.779279 | 0.63173157 | 0.9836963 |
| PVH | 694.75 | 188.081859 | 370.636364 | 129.540938 | 2.07E-05 | 5.48E-05 |
| ARH | 93.375 | 36.1027799 | 17.6363636 | 17.4371599 | 2.27E-06 | 6.63E-06 |
| DMH | 807.75 | 236.467908 | 427.272727 | 178.9425 | 1.83E-05 | 4.86E-05 |
| MEPO | 81.375 | 29.2522893 | 51.1818182 | 13.429953 | 0.00207436 | 0.00462297 |
| MPO | 849.75 | 232.656553 | 283.454545 | 94.7241929 | 1.16E-14 | 7.37E-14 |
| PVp | 130.125 | 54.9634619 | 28.9090909 | 21.9975205 | 1.77E-06 | 5.24E-06 |
| SBPV | 160 | 37.6221819 | 98.4545455 | 35.4185365 | 0.00039098 | 0.00093571 |
| SCH | 27.125 | 11.3821602 | 12.1818182 | 8.65815433 | 0.00574439 | 0.01212593 |
| AHN | 1094 | 245.908229 | 590.727273 | 159.854991 | 2.74E-08 | 9.61E-08 |
| MBO | 805.25 | 216.305175 | 757.545455 | 110.760429 | 0.51172074 | 0.81698157 |
| MPN | 198.5 | 45.033321 | 51.4545455 | 25.8703059 | 5.95E-16 | 4.46E-15 |
| VMH | 153.875 | 38.8529738 | 76.8181818 | 54.2158984 | 0.00247327 | 0.00546123 |
| PH | 1769.25 | 339.613962 | 1866.72727 | 311.243342 | 0.4837325 | 0.7743595 |
| LHA | 5209.625 | 1000.09841 | 3886.90909 | 857.446494 | 0.00140149 | 0.00318858 |
| LPO | 848.125 | 283.128662 | 404.545455 | 109.755513 | 2.10E-08 | 7.47E-08 |
| RCH | 86.25 | 35.2328986 | 32.4545455 | 16.0210089 | 3.64E-06 | 1.03E-05 |
| STN | 367.5 | 74.8484182 | 298.363636 | 56.6926322 | 0.01478468 | 0.02999463 |
| TU | 252.75 | 84.076071 | 147.909091 | 68.1049991 | 0.00417905 | 0.00897972 |
| ZI | 2455.625 | 511.563133 | 4609.18182 | 886.261115 | 4.51E-12 | 2.19E-11 |
| SCs | 4945.75 | 1160.32209 | 3745.09091 | 1042.10983 | 0.01394329 | 0.02843199 |
| IC | 7683.375 | 1588.1039 | 7350.45455 | 1501.29087 | 0.62202223 | 0.96983701 |
| SNr | 1111.375 | 592.97746 | 1656.18182 | 329.724375 | 0.00931997 | 0.01920042 |
| VTA | 1038.625 | 316.671952 | 1671.45455 | 246.471647 | 1.33E-06 | 3.99E-06 |
| RR | 949.75 | 331.713624 | 1543.18182 | 206.262851 | 2.83E-06 | 8.16E-06 |
| MRN | 5083.875 | 1520.67142 | 16277.7273 | 1487.87379 | 7.36E-42 | 5.52E-40 |
| SCm | 7913.375 | 1250.74617 | 9225.18182 | 1469.88318 | 0.0325595 | 0.06420863 |
| PAG | 4731 | 957.865782 | 4070.09091 | 1101.48095 | 0.13674879 | 0.24401484 |
| CUN | 209.25 | 59.4564667 | 298.727273 | 148.495179 | 0.02834945 | 0.05609075 |
| RN | 176.5 | 55.3224315 | 1420.18182 | 113.062654 | 1.12E-130 | 2.70E-128 |
| SNC | 466.375 | 167.249119 | 833.818182 | 183.825362 | 8.98E-07 | 2.73E-06 |
| PPN | 550.375 | 130.480363 | 1371.36364 | 215.255324 | 2.26E-25 | 4.44E-24 |
| RAmb | 929.375 | 173.095298 | 1344.45455 | 296.684467 | 2.42E-05 | 6.31E-05 |
| NLL | 827.5 | 185.683294 | 1539.90909 | 500.970948 | 1.50E-07 | 4.94E-07 |
| PSV | 555.375 | 73.7077966 | 1395.81818 | 630.354633 | 2.67E-10 | 1.11E-09 |
| PB | 719.625 | 218.480426 | 1301.36364 | 859.319297 | 0.00248172 | 0.00546982 |
| SOC | 380.625 | 92.0418189 | 400.545455 | 99.8442423 | 0.63872424 | 0.99329489 |
| B | 52.125 | 29.4833488 | 41.2727273 | 29.6347462 | 0.46034846 | 0.74387844 |
| PCG | 822 | 349.111935 | 1056.18182 | 485.823799 | 0.18381363 | 0.32080428 |
| PG | 3173.5 | 404.568906 | 2755.54545 | 405.212627 | 0.02226692 | 0.04442269 |
| PRNc | 1156.375 | 264.718685 | 4369.27273 | 2403.22779 | 4.39E-15 | 2.96E-14 |
| SUT | 92.625 | 72.2356016 | 208.636364 | 168.71175 | 0.00517259 | 0.01101587 |
| TRN | 2143.375 | 269.771828 | 2479.27273 | 643.581711 | 0.08842236 | 0.16260493 |
| V | 194.875 | 70.5031661 | 606 | 398.200954 | 6.61E-08 | 2.23E-07 |
| CS | 644.25 | 228.900946 | 1433 | 548.215104 | 7.22E-08 | 2.42E-07 |
| LC | 81.375 | 34.0836681 | 65.7272727 | 35.4347031 | 0.3146108 | 0.52831701 |
| LDT | 146.375 | 81.8394552 | 210.454545 | 163.707278 | 0.19968489 | 0.34698867 |
| NI | 179.125 | 84.0415501 | 365.818182 | 204.640572 | 0.00149195 | 0.00338157 |
| PRNr | 2227.75 | 479.209692 | 7805.63636 | 1843.80543 | 2.34E-39 | 1.47E-37 |

**Supplementary Table 4.** Summary region-based statistics table from brain-wide c-fos count comparison between single-housed males exposed to urine and saline (n=4 biologically independent samples for each condition, FDR=0.05 for q-value)

| ROIs | mean.saline | std.saline | mean.urine | std.urine | p-value | q-value |
| --- | --- | --- | --- | --- | --- | --- |
| FRP | 5.75 | 2.21735578 | 10.25 | 11.0867789 | 0.38635903 | 1 |
| MO | 10151 | 2698.73612 | 8798 | 1988.64979 | 0.33405048 | 0.99386234 |
| SS | 44241 | 7294.89995 | 30519.25 | 6498.20931 | 0.00252153 | 0.04651245 |
| ILA | 1400.75 | 205.344223 | 1615.5 | 504.349416 | 0.34562616 | 1 |
| GU | 859.75 | 312.097608 | 732.75 | 257.89969 | 0.49856546 | 1 |
| VISC | 1967.5 | 424.895673 | 1791.75 | 512.350385 | 0.5428334 | 1 |
| AUD | 27404.5 | 2749.26687 | 24041 | 4017.60235 | 0.11740854 | 0.61472856 |
| VIS | 54647 | 5387.75878 | 47429 | 7094.74223 | 0.0638999 | 0.45334898 |
| ACA | 3911.25 | 155.160938 | 3909.25 | 1165.19966 | 0.99708351 | 1 |
| PL | 870 | 161.517801 | 810 | 116.975781 | 0.47636564 | 1 |
| ORB | 995.75 | 270.905119 | 1044 | 433.936247 | 0.82932072 | 1 |
| AI | 3541.25 | 1088.7869 | 3075.75 | 1160.21848 | 0.51048768 | 1 |
| RSP | 15636.5 | 672.227392 | 12938 | 2049.4313 | 0.00834069 | 0.10092931 |
| TEa | 13874 | 2081.0901 | 12132.25 | 1808.16359 | 0.1449849 | 0.67771513 |
| PERI | 5259.75 | 476.1704 | 4620.5 | 742.752314 | 0.11651507 | 0.6141783 |
| ECT | 14308.25 | 1221.11762 | 12866.5 | 2725.90444 | 0.28428824 | 0.90345558 |
| CA | 13124.25 | 649.664721 | 12813.25 | 1860.06584 | 0.7178315 | 1 |
| DG | 5379 | 571.415202 | 5630.75 | 615.147882 | 0.49673518 | 1 |
| ENT | 36884.75 | 1427.92492 | 33744 | 6391.326 | 0.2736083 | 0.89145749 |
| PAR | 3847.75 | 288.674413 | 3848 | 1081.47893 | 0.9996073 | 1 |
| POST | 2881 | 124.362374 | 2468.5 | 298.753522 | 0.00568034 | 0.08108003 |
| PRE | 4709.75 | 516.269552 | 4963 | 553.472071 | 0.45081741 | 1 |
| SUB | 14090.75 | 258.3865 | 12844.5 | 1441.35989 | 0.06676397 | 0.46540698 |
| CLA | 1750.25 | 237.352586 | 1688.75 | 159.616989 | 0.62835086 | 1 |
| EP | 4785 | 770.920662 | 4039.25 | 638.189823 | 0.09876051 | 0.56070126 |
| LA | 1014.75 | 480.082198 | 631.25 | 248.72391 | 0.08923682 | 0.5270687 |
| BLA | 3330.5 | 189.015872 | 2542.75 | 438.212563 | 0.00089385 | 0.02232759 |
| BMA | 2842.25 | 270.991236 | 2227 | 510.866584 | 0.02881308 | 0.26018094 |
| PA | 700 | 87.2505969 | 667.25 | 116.029809 | 0.61919883 | 1 |
| CP | 10138.5 | 1121.0223 | 8415.25 | 1383.27929 | 0.02912966 | 0.26018094 |
| ACB | 1900.5 | 493.296733 | 1662.5 | 308.601685 | 0.36618832 | 1 |
| FS | 265.5 | 54.0709411 | 197.75 | 21.5154983 | 0.00374718 | 0.05933355 |
| OT | 1174.75 | 104.87572 | 783.5 | 107.927445 | 8.41E-09 | 1.12E-06 |
| LS | 2376.5 | 1211.51544 | 2502.75 | 892.922682 | 0.85183751 | 1 |
| SF | 366.5 | 384.349234 | 798 | 196.872209 | 0.21687968 | 0.80839929 |
| AAA | 910.25 | 185.465855 | 587.75 | 179.046316 | 0.00360848 | 0.05846712 |
| CEA | 1589.25 | 264.521423 | 1279.75 | 210.780099 | 0.03846833 | 0.3258291 |
| IA | 512 | 54.4242593 | 432.25 | 104.79305 | 0.15304589 | 0.69656742 |
| MEA | 2974.75 | 481.770606 | 2564 | 550.013333 | 0.20177867 | 0.78295349 |
| GPe | 331.5 | 41.0731056 | 309.75 | 82.1477328 | 0.5755571 | 1 |
| GPI | 109.5 | 12.7148207 | 85.5 | 27.1600196 | 0.07224244 | 0.47507918 |
| SI | 1146.5 | 199.782382 | 798 | 95.7113717 | 0.0001815 | 0.00805979 |
| MA | 222.25 | 90.149413 | 81.25 | 43.0300283 | 0.00079467 | 0.02071317 |
| MSC | 836.25 | 201.769464 | 577.25 | 105.626307 | 0.00309025 | 0.05293153 |
| TRS | 154.25 | 182.540178 | 262.75 | 127.917617 | 0.38043442 | 1 |
| BST | 1766.25 | 305.473812 | 1788.75 | 845.370678 | 0.95588252 | 1 |
| VAL | 114 | 112.519628 | 192 | 175.402395 | 0.33793644 | 0.99799456 |
| VM | 279.75 | 18.1176709 | 265.25 | 142.120547 | 0.82228646 | 1 |
| VP | 184 | 185.854782 | 177.75 | 114.796559 | 0.93890976 | 1 |
| SPF | 515.25 | 72.4310016 | 518 | 138.984412 | 0.96610753 | 1 |
| MG | 785.75 | 150.648321 | 788.5 | 378.314243 | 0.98680155 | 1 |
| LGd | 103 | 112.679486 | 55.25 | 11.8427193 | 0.14238482 | 0.67212363 |
| LP | 400.25 | 453.129397 | 277.5 | 56.7597275 | 0.43269047 | 1 |
| PO | 205 | 271.682413 | 106.5 | 51.6946161 | 0.23342507 | 0.83651208 |
| POL | 114.25 | 17.0171482 | 126.5 | 46.7083147 | 0.53821854 | 1 |
| AV | 352 | 390.560281 | 692.25 | 534.66025 | 0.26369659 | 0.89092653 |
| AM | 592.25 | 419.338666 | 756.75 | 452.259789 | 0.57840833 | 1 |
| AD | 161 | 185.817114 | 272 | 109.69959 | 0.35633741 | 1 |
| LD | 126 | 91.6587875 | 171.25 | 80.4751514 | 0.4368459 | 1 |
| IMD | 304 | 179.244712 | 308 | 67.6214956 | 0.96288123 | 1 |
| MD | 660.75 | 418.517523 | 885.25 | 220.283719 | 0.35233694 | 1 |
| SMT | 21.75 | 13.4008706 | 40.25 | 33.1197323 | 0.20602717 | 0.7892223 |
| PVT | 2051.25 | 896.739046 | 2448.25 | 263.596124 | 0.38897174 | 1 |
| PT | 458 | 220.215046 | 585.25 | 197.292296 | 0.34093556 | 1 |
| RE | 1503.5 | 546.468968 | 1219.5 | 522.563234 | 0.43931411 | 1 |
| CM | 447.75 | 118.015183 | 447 | 131.929274 | 0.99240185 | 1 |
| PCN | 48 | 36.4508802 | 63.25 | 38.7416658 | 0.52625516 | 1 |
| CL | 138.25 | 114.59603 | 160.25 | 49.614346 | 0.73178738 | 1 |
| PF | 294.75 | 234.362077 | 342 | 136.342705 | 0.72124626 | 1 |
| RT | 539 | 202.808613 | 393 | 129.282636 | 0.14065112 | 0.66950901 |
| LGv | 671.75 | 81.6797609 | 686 | 150.890689 | 0.85185968 | 1 |
| MH | 99.25 | 35.0273703 | 144.75 | 48.1412851 | 0.10495892 | 0.58262641 |
| LH | 247.25 | 132.77142 | 288.25 | 79.5628682 | 0.56701022 | 1 |
| PVH | 744.75 | 175.262422 | 644.75 | 212.503137 | 0.44481552 | 1 |
| ARH | 89.75 | 12.3119183 | 97 | 53.4290807 | 0.77934917 | 1 |
| DMH | 922 | 227.190082 | 693.5 | 209.880125 | 0.11077879 | 0.60661487 |
| MEPO | 83.25 | 22.0056811 | 79.5 | 38.7685439 | 0.86500543 | 1 |
| MPO | 1017.5 | 176.299934 | 682 | 142.05398 | 0.00087888 | 0.02232759 |
| PVp | 157.5 | 53.2697538 | 102.75 | 47.0416482 | 0.111932 | 0.60665249 |
| SBPV | 187.5 | 22.4276615 | 132.5 | 27.9821372 | 0.00059798 | 0.01707085 |
| SCH | 32.5 | 12.2338329 | 21.75 | 8.69386757 | 0.10507286 | 0.58262641 |
| AHN | 1298.5 | 161.50645 | 889.5 | 59.1072472 | 1.47E-09 | 2.93E-07 |
| MBO | 919 | 121.81133 | 691.5 | 244.601036 | 0.07097627 | 0.47507918 |
| MPN | 224.75 | 38.6382798 | 172.25 | 37.4377261 | 0.02864844 | 0.26018094 |
| VMH | 161.5 | 33.1511689 | 146.25 | 47.6261483 | 0.57004251 | 1 |
| PH | 2010 | 240.570156 | 1528.5 | 238.087519 | 0.00155437 | 0.03328014 |
| LHA | 6075.5 | 572.94706 | 4343.75 | 78.8601082 | 2.02E-15 | 1.21E-12 |
| LPO | 1060 | 236.455775 | 636.25 | 106.881165 | 3.14E-05 | 0.0022162 |
| RCH | 103.25 | 37.9857429 | 69.25 | 26.1326743 | 0.0834592 | 0.50335405 |
| STN | 418.25 | 67.5394946 | 316.75 | 40.5246838 | 0.00237906 | 0.04527774 |
| TU | 307.25 | 68.4367104 | 198.25 | 62.3665241 | 0.00776885 | 0.09806855 |
| ZI | 2745.75 | 433.43925 | 2165.5 | 445.304765 | 0.03886035 | 0.3258291 |
| SCs | 5552.75 | 963.354685 | 4338.75 | 1109.45674 | 0.05834102 | 0.4317956 |
| IC | 8121 | 2181.89871 | 7245.75 | 783.229798 | 0.34457464 | 1 |
| SNr | 1506.5 | 614.419238 | 716.25 | 163.128528 | 0.00022185 | 0.00859915 |
| VTA | 1233 | 298.937006 | 844.25 | 209.463402 | 0.01140145 | 0.12896544 |
| RR | 977 | 482.340129 | 922.5 | 148.706646 | 0.79091902 | 1 |
| MRN | 4564.25 | 1032.19616 | 5603.5 | 1900.06342 | 0.23652276 | 0.839026 |
| SCm | 7331 | 952.152299 | 8495.75 | 1356.15815 | 0.09407589 | 0.54001546 |
| PAG | 4143.25 | 569.651575 | 5318.75 | 946.121689 | 0.00785651 | 0.09812458 |
| CUN | 197.5 | 57.2625532 | 221 | 67.8331286 | 0.53220721 | 1 |
| RN | 176 | 42.4735212 | 177 | 73.0524925 | 0.97773488 | 1 |
| SNC | 571.75 | 187.297224 | 361 | 24.0416306 | 0.00091366 | 0.02235674 |
| PPN | 536 | 171.388448 | 564.75 | 98.996212 | 0.7518104 | 1 |
| RAmb | 1009.25 | 209.281907 | 849.5 | 95.3886786 | 0.07982648 | 0.49083053 |
| NLL | 912.5 | 222.982062 | 742.5 | 107.059174 | 0.09198063 | 0.53797454 |
| PSV | 599.75 | 36.6731055 | 511 | 77.9786296 | 0.02236944 | 0.21986427 |
| PB | 722 | 260.435533 | 717.25 | 208.655018 | 0.97335098 | 1 |
| SOC | 369.5 | 91.7769034 | 391.75 | 104.948797 | 0.7152767 | 1 |
| B | 36.5 | 26.4007576 | 67.75 | 26.0816027 | 0.06562305 | 0.46012885 |
| PCG | 669.25 | 345.709295 | 974.75 | 320.389737 | 0.1167912 | 0.6141783 |
| PG | 3352.75 | 191.102721 | 2994.25 | 509.616441 | 0.16537059 | 0.71323501 |
| PRNc | 1195 | 329.24459 | 1117.75 | 226.11999 | 0.65902196 | 1 |
| SUT | 125.25 | 89.9865731 | 60 | 35.2041664 | 0.06835258 | 0.46565198 |
| TRN | 2259 | 320.609836 | 2027.75 | 177.118369 | 0.13814973 | 0.66317109 |
| V | 218.75 | 90.0902325 | 171 | 44.2944692 | 0.26576754 | 0.89092653 |
| CS | 698.75 | 326.305966 | 589.75 | 88.658051 | 0.40341985 | 1 |
| LC | 71.5 | 26.4133804 | 91.25 | 41.8678476 | 0.305915 | 0.94049253 |
| LDT | 129.75 | 113.793307 | 163 | 44.0681291 | 0.53475862 | 1 |
| NI | 169 | 87.5861481 | 189.25 | 92.3882207 | 0.7047483 | 1 |
| PRNr | 2518.5 | 461.991703 | 1937 | 311.394712 | 0.0171586 | 0.17735484 |

